# Norepinephrine as a Spatial Memory Reset Signal

**DOI:** 10.1101/2020.06.07.138859

**Authors:** Stephanie L. Grella, Sarah M. Gomes, Rachel E. Lackie, Briana Renda, Diano F. Marrone

**Author notes:** Corresponding & First Author: Stephanie L. Grella, Ph.D., Address: Department of Psychology, Wilfrid Laurier University, 75 University Ave, Waterloo, ON, N2L 3C5, Canada. CONFLICT OF INTEREST The authors declare no competing financial interests.

## Abstract

Contextual information is represented in the hippocampus (HPC) partially through the recruitment of distinct neuronal ensembles. It is believed that reactivation of these ensembles underlies memory retrieval processes. Recently, we showed that norepinephrine (NE) input from phasic locus coeruleus (LC) activation induces hippocampal plasticity resulting in the recruitment of new neurons and a disengagement from previously established representations. We hypothesize that NE may provide a neuromodulatory, mnemonic switch signaling the HPC to move from a state of retrieval to encoding in the presence of novelty, and therefore, plays a role in memory updating. Here, we tested whether bilateral dorsal dentate gyrus (DG) infusions of the β-adrenergic receptor (BAR) agonist isoproterenol (ISO), administered prior to encoding or retrieval, would impair spatial working and reference memory by reverting the system to encoding (thereby recruiting new neurons) potentially interfering with retrieval of the previously established spatial ensemble. We also investigated whether dDG infusions of ISO could promote cognitive flexibility by switching the system to encoding when it is adaptive (i.e. when new information is presented e.g. reversal learning). We found that intra-dDG infusions of ISO given prior to retrieval caused deficits in working and reference memory which was blocked by pre-treatment with the BAR-antagonist, propranolol (PRO). In contrast, ISO administered prior to reversal learning led to improved performance. These data support our hypothesis that NE serves as a novelty signal to update HPC contextual representations via BAR activation-facilitated recruitment of new neurons. This can be both maladaptive and adaptive depending on the situation.

**SIGNIFICANCE STATEMENT:** The current work highlights the involvement of hippocampal BARs in determining the flexibility of contextual representations to promote new learning in a way that supports adaptive behavior. This work builds upon previous work showing that noradrenergic input to the hippocampus is involved in recruiting new neurons resulting in new contextual representations and may be involved in the underlying neural mechanisms that support memory updating. These data suggest targets for anxiety disorders such as PTSD, which are characterized by noradrenergic dysregulation, and may also involve impairments in memory updating mechanisms where the incorporation of new information is not effectively encoded. The further understanding of the neurobiological mechanisms involved in updating memories may provide insight into novel treatment strategies.

## INTRODUCTION

Recruitment of new hippocampal neurons is part of memory encoding where active neuronal ensembles form contextual representations of discrete experiential episodes. Tasks involving retrieval require reactivation of the representations formed during encoding (Guzowski et al., 1999; Chawla et al., 2005; Garner et al., 2012; Pevzner et al., 2012; Josselyn et al., 2015; Tonegawa et al., 2015a, 2015b; Eichenbaum, 2016). If those representations remapped (i.e., a new cellular ensemble was recruited, rather than reactivation of the cells comprising the previously formed representation) this should theoretically result in a retrieval error. Therefore, switching the memory system back to a state of encoding would prove maladaptive in situations where retrieval is necessary to perform a memory task. Unless new information was at hand. In this case, it would be adaptive to incorporate that new information into the memory trace and theoretically retrieval and encoding would occur together to update that representation.

Furthermore, it is hypothesized that post-traumatic stress disorder (PTSD) involves impairments in memory-updating mechanisms where incorporation of new information (e.g., safety signals) is not encoded at a functional level which may reflect an inability to remap hippocampal contextual representations (i.e., trauma-related representations are reactivated rather than incorporating safety signals into existing memory traces) (Maren et al., 2013; Morrison and Ressler, 2014; Giustino et al., 2016; Liberzon and Abelson, 2016; Elsey and Kindt, 2017; Lee et al., 2017; Sheynin and Liberzon, 2017). The pathophysiology of anxiety disorders such as PTSD is characterized by noradrenergic dysregulation (Hendrickson and Raskind, 2016). We believe the locus coeruleus (LC), the site of noradrenergic cell bodies, plays a role in memory updating, specifically by biasing the system towards encoding. This suggests that individuals with PTSD may be experiencing an inability to engage this transition to encoding through dysfunction of the LC norepinephrine (NE) system.

The LC responds to salience cues including novelty and sends a major noradrenergic projection to the dentate gyrus (DG) (Aston-Jones and Bloom, 1981; Vankov et al., 1995; Berridge and Waterhouse, 2003; Harley, 2007; Aston-Jones and Waterhouse, 2016). LC activation causes NE release (Blackstad et al., 1967; Fuxe et al., 1968; Ungerstedt, 1971; Lindvall and Björklund, 1974; Pickel et al., 1974; Ross and Reis, 1974; Babstock and Harley, 1992; Frizzell and Harley, 1994) and induces downstream hippocampal plasticity (Bliss et al., 1983; Neuman and Harley, 1983; Stanton and Sarvey, 1985; Lacaille and Harley, 1985; Gray and Johnston, 1987; Hopkins and Johnston, 1988; Harley, 1991; Klukowski and Harley, 1994; Walling and Harley, 2004; Almaguer-Melian et al., 2005; Lashgari et al., 2008; Lemon et al., 2009; Lim et al., 2010; Walling et al., 2011; Hagena et al., 2016), effects which are β-adrenergic receptor (BAR)-dependent (Kitchigina et al., 1997). It is thought that LC-NE activation induces changes in network dynamics occurring at critical times when learning is necessary to promote adaptive behavior e.g., reversal learning (Sara et al., 1994). These configurations function to reset the system (Bouret and Sara, 2005) as novelty-associated activation of the LC causes the hippocampus (HPC) to recruit a new population of neurons to represent the immediate context (i.e., global remapping) (Grella et al., 2019). This observation is consistent with the idea that NE provides a reset signal promoting global remapping in the HPC in the presence of new information and our hypothesis that it can bias memory towards encoding. This hypothesis suggests that the effect of modulating NE on memory will depend on the stage of training.

Although NE has been shown to mediate different stages of memory (McGaugh et al., 1990; Do Monte et al., 2008) it is unclear how it is involved in updating memory. To assess this, we investigated how activation and blockade of BARs exerts modulatory influence during learning and recall. We tested whether infusions of the BAR-agonist isoproterenol (ISO) would impair working and reference memory retrieval by switching the system to encoding when it is potentially maladaptive (e.g., when retrieval of a previously established ensemble is required to complete a task). Given that LC neurons exhibit plasticity as a function of environmental contingency changes to promote adaptive behavior, (Sara and Segal, 1991) we also tested whether ISO would, in contrast, enhance cognitive flexibility by promoting encoding when it is potentially adaptive (e.g., when new information is presented) and whether these effects could be blocked with the BAR-antagonist propranolol (PRO).

## MATERIALS AND METHODS

### Animals

Eighty-seven adult male Fischer-344 rats were purchased from Harlan (Indianapolis, IN) at 16 weeks old (∼350-375g) for experiment 1 & 2 and at 10 weeks old (∼300-325g) for experiment 3. The number of rats included in each experiment and group are listed in **Table S1** and in the figure legends. Rats were housed in standard transparent Plexiglas cages (47.6cm L x 26.0cm W x 20.3cm H), pair-housed initially and then single-housed after surgery. They were kept on a 12:12 hour reverse light cycle (lights ON at 7pm) and provided with food and water *ad libitum* until they recovered from surgery after which the animals in experiment 1 were food restricted to 90% of their free fed weight. Animals in experiments 2 & 3 remained on an *ad libitum* diet. All procedures were approved by the Wilfrid Laurier University Animal Care Committee in accordance with the guidelines of the Canadian Council on Animal Care.

### Surgery

For 4 consecutive days prior to surgery animals were weighed, handled for 15 min, and given 20 g of a nutritionally complete dietary supplement containing trimethoprim / sulfamethoxazole antibiotic (MediGel® TMS; ClearH20, Westbrook, ME) in addition to their regular diet in their home cage. The following day, rats underwent implantation of a bilateral guide cannula. Several days prior to surgery, two 22-gauge stainless steel guide cannulas (Plastics One, Roanoke, VA) were cemented together to form a bilateral cannula and left to dry. The next day they were autoclaved and again left to dry. At the start of surgery, rats were deeply anesthetized with 5% isoflurane and 70% oxygen, (induction) and maintained at a level of 2-3% isoflurane for the duration of the surgery. They were anchored in a stereotaxic frame with ear bars to ensure a flat skull surface and prepped for aseptic surgery. Rats were administered a sub-cutaneous (s.c.) injection of ketoprofen (Anafen®; Sigma Aldrich, Oakville ON; 0.15 ml of a 10 mg/mL solution) for general analgesia, and 3ml of sterile physiological saline (0.9% NaCl; s.c.) for fluid replacement in case of blood loss. A midline incision was made on the scalp and 6 holes were drilled. Each rat was implanted with the bilateral cannula (8mm in length, Plastics One) aimed at the dorsal DG with the coordinates: AP -3.3mm, ML +/- 2.1mm, DV -4.2mm (from skull) relative to Bregma (Paxinos and Watson, 2013). Cannulae were anchored to the skull with four skull screws (#0-80, Plastics One) and dental acrylic. At the end of surgery stainless steel stylets (flush with guide) were screwed into the cannulae to ensure patency and rats were placed on a heating pad for 1 h. They were given an additional 0.15 ml injection (s.c.) of ketoprofen (Sigma Aldrich, Oakville, ON) 24 h later and allowed 7 d for recovery undisturbed except for daily weighing. During the first 4 days of recovery rats continued to receive 20 g of TMS in their home cage and given their regular diet mixed with water in mashed form in addition to regular chow pellets.

### Drugs and Infusions

Rats received either (-)-isoproterenol bitartrate [ISO; 10ug/ul (5ug/side) dissolved in sterile saline; Sigma Aldrich, Oakville, ON] or (+/-) -propranolol hydrochloride (DL) [PRO; 3ug/ul (2.5ug/side) dissolved in sterile saline; Sigma Aldrich]. Given that few experiments have targeted the DG with these specific drugs in awake, freely moving animals, the doses we chose were based on a literature search **(Table S2).** We decided to infuse 5ug in the DG of each hemisphere since Geyer and Masten (1989) found that infusion of a similar amount resulted in an increase in diversive (reconnaissance-like) exploration. For PRO, no studies had targeted the DG specifically. Ji et al. (2003) and Chai et al. (2014) found impaired memory consolidation and a blockade of NE-facilitated memory enhancements when they targeted the CA1 (5ug per side). However, Hatfield and McGaugh (1999) and Barsegyan et al. (2014) found spatial memory impairments and diminished NE-facilitated memory enhancements with smaller amounts (0.3, and 1ug) when targeting the BLA. Therefore, we decided to use 1.5 μg for PRO. For each infusion, stylets were unscrewed from each rat’s cannulae and a 30-gauge infusion cannula (1mm below pedestal) connected via polyethylene tubing (PE-10) to a 10-μL Hamilton syringe mounted onto a microfluidic infusion pump (Harvard Apparatus, model: 70-2000) was inserted into the guide cannulae. Rats were infused with 0.5μL on either side of the brain at a rate of 0.5μL/min and the infusion cannula left in place for 1 min post-infusion to ensure the liquid had diffused from the injection site.

## EXPERIMENT 1: DELAYED NON-MATCH TO POSITION (DNMP)

### Apparatus

We used a radial arm maze (122cm in diameter; Stoelting Co., Wood Dale, IL), which consisted of 12 grey, equidistantly spaced, polyethylene arms (50cm L x 10cm W x 13cm H) that radiated from a small circular rotating central platform. The maze rested on a table, elevated 84cm from the ground, located in the center of the room (2.44m L x 2.24m W x 2.95m H) and extra-visual cues (geometric shapes) were positioned on the walls. Other visual cues included a computer in one corner of the room.

### Pilot Experiment

To decide which arm separations to use for the task, we ran a pilot study with animals that did not undergo surgery. These rats received habituation and pre-training trials and then 6 acquisition-training sessions (see below). Each acquisition-training session consisted of 6 trials (sample + choice) per day to assess performance on arm separations 1-6 (order counterbalanced). We measured latency to obtain the reward, the number of errors made and the percentage of trials where a correct response was made for the last 3 days.

### Procedure

The DNMP task consisted of 5 stages: 1) Habituation 2) Pre-training 3) Acquisition 4) Testing 5) Curtain probe test. Timeline and task schematic outlined in Figure 1A-C.

**Figure 1.**
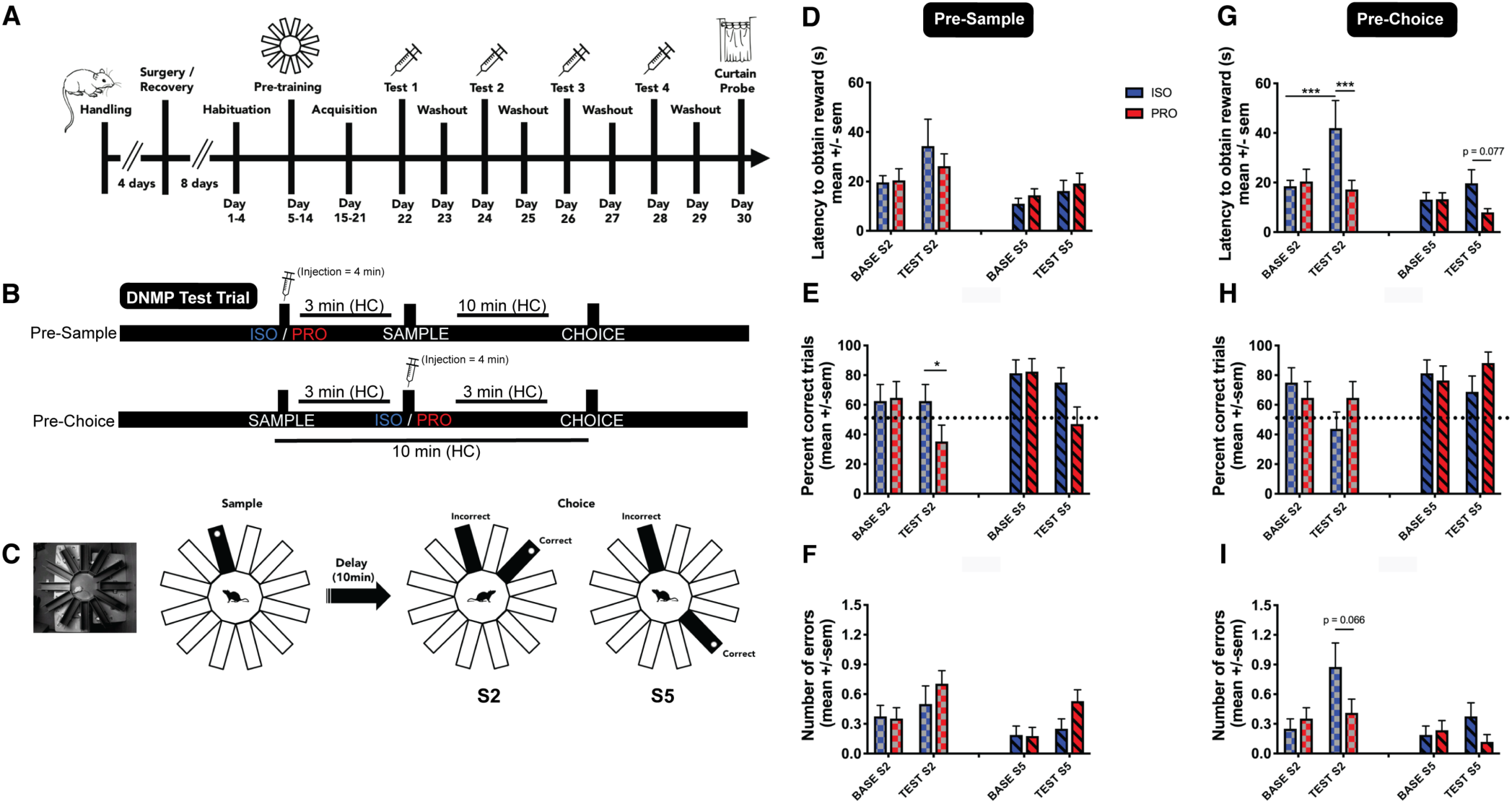
(A) Experiment 1 timeline. (B) Timeline for infusions on test days. (C) Schematic showing DNMP task. During the sample phase, one arm is open and baited and animals are trained to obtain a reward from this arm. After a delay of 10 min, in the choice phase, animals are placed back in the maze and presented with a choice between the previously rewarded arm and the new arm. The sample and choice arms are separated by either 2 (S2) or 5 (S5) arms making the task DG-dependent or DG-independent respectively. (D-I) Test Day - animals were tested on 4 conditions on 4 different days with each test day separated by a washout day. Rats were assigned to either the ISO (blue; n = 20) or PRO (red; n = 20) group and received a drug infusion either 3 min prior to the sample phase (Pre-Sample) or the choice phase (Pre-Choice) and were tested in the S2 (checkered) or S5 (striped) condition. TRIAL 1: habituation trial (not shown); TRIAL 2: baseline trial (BASE) - animals infused with saline; TRIAL 3: test trial (TEST) - animals received drug treatment they were assigned to. Significant differences denoted with asterisks * p < 0.05, **p < 0.01, ***p < 0.001. DNMP = Delayed non-match to position, DG = Dentate gyrus, HC = Home cage, ISO = Isoproterenol, PRO = Propranolol, S2 = two arm separation, S5 = 5 arm separation, PS = Pre-sample, PC = Pre-choice.

### Habituation & Pre-Training

Habituation lasted 4 days. On day 1, rats were given one 20-min habituation trial where they freely explored the maze. All 12 arms were open and baited with a reward placed in a small plastic grey cup at the end of the arm. The next day rats were given two 10 min trials (inter-trial interval = 1 h) in the maze with 6 arms open and baited. For the next two days, they were given two 5 min trials a day with 3 arms open and baited. On day 5 rats began Pre-training. Pre-training lasted 10 days. During this phase rats were given two trials / day with only one arm open and baited. The goal was to train the rats to retrieve the reward in less than 2 min. By the tenth day all rats could do this. On day 14, rats began acquisition training.

### Acquisition Training

Animals received 4 trials / day. Each trial consisted of 2 phases: sample & choice, separated by a 10-min delay. During the sample, all arms except the sample arm were blocked off. The rat was placed in the center of the maze and permitted to visit the sample arm and obtain half a Froot Loop®. Latency to obtain the reward was recorded. Once the animal retrieved the reward, he was left in the maze an additional 10 s to promote memory for the sample arm location. The rat was then placed back in his home cage (HC) and 10 min later tested on the choice phase. During the 10 min delay, the maze was rotated, preserving arm-location, yet eliminating the possibility of odor being used as an intra-maze cue. During the choice phase, the previously rewarded sample arm was now unrewarded. An additional correct arm was open and rewarded. Correct arms varied in distance from the sample arm by a spatial separation of 2 (S2), or 5 (S5) arms **(Figures 1B & C)**. Cups in each of the two arms appeared identical from afar and both contained half a Froot Loop®, but the cup in the unrewarded arm had a mesh overlay preventing access to the reward. Latency to reach the reward, choice accuracy, and number of errors were recorded. When the incorrect arm was chosen, the rat was permitted to self-correct. Re-entry into the incorrect arm was considered an additional error. When a correct choice was made, an additional full Froot Loop® was given in the HC immediately after the trial ended. Rats were given 4 trials (sample + choice) per day (2x S2 & 2x S5) of pseudo-randomly presented combinations of sample + correct arms **(Tables S3 & S4)** [inter-trial interval (ITI) = 90 min until a criterion of 4/6 correct choices were made on S5 trials across 3 consecutive days]. Criterion reached within 6-7 days.

One hour after the last acquisition training trial, stylets were unscrewed from each rat’s cannulae, and the infusion cannula was inserted to make sure the cannula was not blocked. The infuser was left in the cannula for 2 min on each side of the brain to simulate what would occur during testing, but no fluid was delivered. This was done in attempts to reduce the elicitation of a nonspecific stress response on test day. Following this, the dust caps were screwed back in and animals were then returned to their HC.

### Test Day

Using a balanced Latin Square design **(Table S5),** animals were tested on 4 conditions on 4 different days with each test day separated by a one-day washout **(Figure 1A).** We used a 2x2x2 design with a between-subject factor of GROUP (drug treatment), a within-subject factor of INFUSION TIME, and a within-subject factor of ARM-SEPARATION. Rats were assigned to a drug treatment following acquisition (ISO or PRO). This remained constant throughout testing. On test day, rats were infused 3 min prior to either the sample phase (Pre-Sample, PS) or the choice phase (Pre-Choice, PC) and were tested on S2 and S5 conditions. So, all animals were tested on PS-S2, PS-S5, PC-S2, and PC-S5. On each of the 4 test days, instead of receiving two S2 and two S5 trials (as in training), animals received all 4 trials in the condition they were being tested (all S2 or all S5) allowing TRIAL to be included as an additional within-subject factor making a 2x2x2x3 design. Trial 1: Habituation - like the previous day, infusion cannulae were inserted but no fluid was infused; Trial 2: Baseline - animals were infused with sterile saline (0.9% NaCl); Trial 3: Test - animals received either ISO or PRO. In our analyses, we compared Baseline to Test (see **Table S6** for all conditions). During testing, the latency to obtain the reward, the number of errors, and the percent correct trials were measured. The next day after the final test day, rats were given one more washout and the following day, a curtain probe test.

### Washout Sessions and Curtain Probe

Between each test day, animals were given a washout day that was identical to acquisition training to allow the drug to clear before recommencing testing. To ensure that animals were using extra-maze cues rather than intra-maze or interoceptive cues to complete the DNMP task, following the last washout day animals were given a curtain probe. The procedure for both the washout sessions and the curtain probe were identical to acquisition training. For the curtain probe, the exception was that a blue curtain was hung from the ceiling in a circular fashion, surrounding the maze such that animals could not see any of the cues in the room except for the webcam above and a partial view of a few ceiling tiles.

## EXPERIMENT 2: ELEVATED PLUS MAZE (EPM)

### Apparatus

To assess the effects of ISO and PRO on locomotion and anxiety, a separate group of rats were tested using a grey polyethylene EPM (Stoelting Co., Wood Dale, IL) consisting of two open and two closed runways (50cm L x 10cm W x 40cm H) elevated 40cm from the ground. EPM testing took place in a smaller room (1.83m L x 1.78m W x 2.95m H) where the maze was positioned in the center. A separate group of animals was used since there were no drug-naïve animals in the DNMP experiment to serve as the vehicle group for EPM testing.

### Procedure

Rats underwent similar handling and surgical procedures as above. Following recovery, rats were split into 3 groups: ISO, PRO, and vehicle. Using the same doses as above, rats were given a bilateral intra-DG infusion of either ISO, PRO, or vehicle and then 3 min later tested in the EPM. Rats were placed at the junction of the four arms at the beginning of the session. Their behavior was monitored for 5 min. Anxiety-like behavior was assessed by measuring the percentage of time spent in the open arms of the maze compared to the closed arms and the number of entries into the open and closed arms. General locomotor activity was assessed by measuring total number of arm entries.

## EXPERIMENT 3: BARNES MAZE

### Apparatus

The Barnes maze (Barnes, 1979) consisted of a grey circular polyethylene disk (122cm diameter) with 20 circular equidistant holes (10cm diameter; 9.65cm between holes) located around the perimeter of the maze (1.3cm from the edge). The maze was elevated 90cm from ground and beneath each hole was a slot where an escape box (35.56 L x 13.34cm W x 10.16cm H) or a “false” escape box (11.43cm L x 13.34cm W x1.9cm H) could be inserted. For any given trial 19/20 holes were connected to a false escape box and only 1-hole lead to the true escape box. The false escape boxes were significantly smaller than the true escape box; therefore, rats could not escape the maze via these boxes. Their main purpose was to conceal any visual cues that may be apparent from a distance or through an open hole. Four bright white lights (150W) were mounted above the maze, which illuminated the entire maze area. The rest of the room was dark when testing. Animals were motivated to escape from the brightly lit, open platform into the dark, recessed escape box due to their natural tendency to seek out dark, closed spaces. The maze was in the center of a large room (4.5m L x 3.35m W x 2.95m H) and extra-maze visual cues (geometric shapes) were positioned on the walls. Other visual cues included several desks and cabinets.

### Procedure

The experiment consisted of 9 distinct phases: 1) Habituation 2) Acquisition training 3) Acquisition probe test 4) Re-training 5) Curtain probe test 6) Second re-training 7) Reversal training day 1 8) Reversal training days 2-5 and 9) Reversal probe test **(Exp. timeline and task schematic Figures 3A & 3B).**

**Figure 3.**
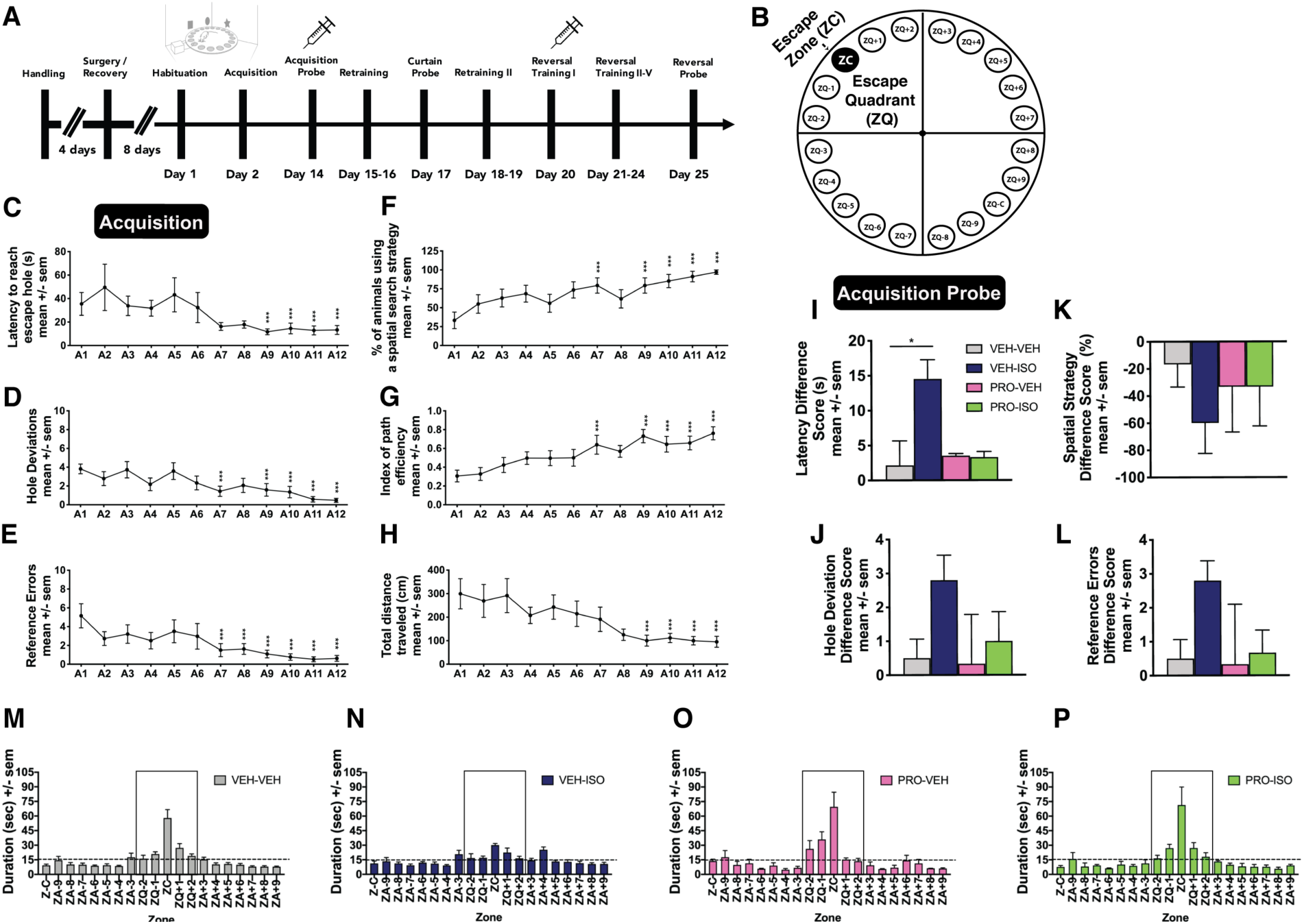
(A) Experiment 3 timeline. (B) Schematic showing the zones of the Barnes maze. The escape hole (black filled in circle ZC) is where the escape box was located. The two holes to the left (ZQ-1, ZQ-2) as well as the two holes to the right (ZQ+1, ZQ+2) contained a false escape box that the rats could not escape into but looked similar to the rats. These five holes comprised the escape quadrant (ZQ). During acquisition training rats learned the spatial location of the escape box, which was consistent across trials. We measured C) latency to reach the escape hole, D) hole deviations E) reference errors F) the percentage of rats using a spatial search strategy, G) path efficiency, and H) the total distance traveled. Animals were then given a 5-min acquisition probe test where the escape box was removed and replaced with a false escape box. Fifteen min prior to the test, rats were given an infusion of either saline or PRO and then placed back in their HC. Seven min prior to the test rats were given another infusion of either saline or ISO. Groups: VEH-VEH (grey; n = 6), VEH-ISO (blue; n = 5), PRO-VEH (pink; n = 3), and PRO-ISO (green; n = 3). We compared performance on the last day of acquisition to performance during the acquisition probe across groups using a difference score for (I) latency, (J) hole deviations, (K) percentage of animals using a spatial search strategy, and (L) reference errors. (M-P) During the acquisition probe, the maze was divided into 20 equal zones. Time spent in the escape zone and quadrant were calculated. Chance levels are expressed as a dotted line. Significant differences denoted with asterisks * p < 0.05, **p < 0.01, ***p < 0.001. ISO = Isoproterenol, PRO = Propranolol, VEH = Vehicle (saline), ZC = Escape zone, ZQ = Escape quadrant.

### Habituation

Rats were given one 5 min habituation trial where they freely explored the maze and could descend into the escape box. Once they entered the escape box, they were permitted to stay in the box for 30 s and were then removed and placed back into the center of the maze until the end of the 5 min period.

### Acquisition Training

Each trial (except habituation) began with a 5 s acclimatization period during which the rat was being held in the start box in the center of the maze. Trials began automatically after the 5 s delay and the start box was lifted. For the first 4 days (A1-A4), rats were given 3 trials / day and during the following 8 days (A5-A12) this was reduced to 2 trials / day for a total of 28 trials with an inter-trial-interval (ITI) of 2 hours on all days.

Including the habituation trial animals received a total of 29 trials prior to the Acquisition Probe Test. Animals learned the spatial location of the escape box, which stayed consistent. Each trial lasted up to 5 min during, which, ANY-maze software recorded the latency to reach the escape hole, total distance travelled, and path efficiency (see below for description). The experimenter recorded the number of reference errors the animal made prior to reaching the escape hole, the number of hole-deviations there were between the first hole visited and the escape hole, and the search strategy used to find the escape hole. If the rat did not find the escape hole in the time allotted, it was gently guided to the escape box. Once the rat was inside, it remained there for 30 s before returning to its HC. For all training trials, rats were grouped into squads of 3-4 where all members of a squad completed a given trial before subsequent trials were run.

### Cardinal Direction at Start

Before testing began rats were pseudo-randomly assigned to one of 4 possible escape locations. These locations were equidistant positioned at 90-degree intervals (North, West, South, East). This was to prevent odor cues from becoming saturated around any one hole, although the maze was cleaned with 10% ethanol between trials to eliminate any odors. Given that each rat was placed in a holding box for a 5 s acclimatization period at the start of each trial, we could not choose the direction the rat would be facing when the trial began. To ensure that this was counterbalanced for north, west, south and east directions, the videos were scored (n = 850) by a researcher blind to the conditions of the experiment. The results are listed in **Table S7**. This was necessary to assess whether rats were using a fixed motor response to find the escape hole.

### Measures

To record behavior in all three testing rooms, a webcam connected to a computer running ANY-maze tracking software (Stoelting Co., Wood Dale, IL) was mounted above each apparatus on the ceiling and behavior was tracked using ANY-maze software (Stoelting Co., Wood Dale, IL).

*Path efficiency* is represented as an index of the efficiency of the path taken by the rat to get from the first position in the test (start) to the last position (escape hole). A value of 1 is indicative of perfect efficiency (e.g., the animal moved in a straight line from the start to the escape hole). It is calculated by dividing the straight-line distance between the first and the last position by the total distance traveled by the rat. This measure was not used during probe sessions, as it cannot be analyzed across time.

*Reference errors* were recorded as a rat dipping its head into any hole other than the escape hole. Repeated dips into the same hole were considered a single error.

*Hole deviations* were quantified as the number of escape holes between the true escape hole and the location in which the animal’s head, first entered a false escape hole. This ranged between 0-10.

There were three possible search strategies: (1) *Random (RD)* – this occurred when the animal moved about the maze in a random, un-systematic manner, searching the same hole more than once and moving into the center of the maze often. (2) *Serial (SE)* – Animals that used a serial search strategy first visited a hole more than two-hole deviations away from the escape hole and then in a serial fashion systematically checked adjacent holes until reaching the escape hole. The animals search path was classified as serial even if he did not make any errors but visited a location at the edge of the maze more than two holes away. (3) *Spatial (SP)* - This search strategy occurred when a rat moved directly from the center of the maze to the correct escape hole or any hole within two-hole deviations away on either the left or right side of the escape hole.

One hour after the last acquisition training trial, stylets were unscrewed from each rat’s cannulae, and the infusion cannula was inserted to make sure the cannula was not blocked. The infuser was left in the cannula for 2 min on each side of the brain to simulate what would occur during testing, but no fluid was delivered. This was done in attempts to reduce the elicitation of a nonspecific stress response during the acquisition probe test. Following this, the stylets were screwed back in and the animals were returned to their home cage.

### Acquisition Probe

Rats were given a 5 min acquisition probe where the escape box was removed and replaced with a false escape box. The maze was rotated to ensure the animals were using extra-maze visuospatial cues to find the escape hole. The maze was divided into 20 equal zones. The escape zone (ZC) contained the escape hole, and the escape quadrant (ZQ) contained the escape zone plus the 2 zones to the left and right (ZQ-2, ZQ-1, ZQ+1, ZQ+2) **(Figure 3B).** Time spent in the escape zone and quadrant were calculated as well as latency to reach the escape hole, number of reference errors, hole deviations, spatial strategy used, and distance traveled.

Fifteen min prior to the test, rats were infused with either saline (VEH) or PRO and returned to their HC. Seven min prior to the test rats were given another infusion of either VEH or ISO. Infusion volume, rate, and procedure were the same as the previous infusion. Rats were then placed back in the HC and tested 3 min later (each infusion took 4 min). This resulted in 4 groups: VEH-VEH, VEH-ISO, PRO-VEH, and PRO-ISO. Following the acquisition probe animals received 2 days of retraining (2 trials per day, ITI = 2 hours) to reduce any extinction learning that may have occurred during the acquisition probe (probe fixed in length; no escape box).

### Curtain Probe

Animals were re-trained following the acquisition probe and then given a curtain probe trial. The purpose of the curtain probe was to assess whether rats were using intra or extra-maze cues to locate the escape box. The procedure for this test was identical to the acquisition probe except that animals did not receive any infusions and a brown plastic curtain was hung around the maze from the ceiling effectively blocking all visual access to the rest of the room. After this test animals received an additional 2 days of retraining (2 trials per day; ITI = 2h) to reduce any extinction learning that may have occurred during the curtain probe trial.

### Reversal Training

Similar to the acquisition probe, 1 hour after the last retraining trial, stylets were unscrewed from each rat’s cannulae, and the infusion cannula was inserted and left in the guide cannula for 2 min on each side of the brain, stylets were then screwed back in and animals were returned to their HC. The following day animals received their first reversal training trial.

Reversal training was similar to acquisition training except the location of the escape box was moved 180 degrees. It lasted 5 days with one trial on the first day and 2 trials / day (ITI = 2 hours) after that. Fifteen min prior to the first reversal training trial rats were given an infusion of either VEH or PRO. Seven min prior to RV1 rats were given another infusion of either VEH or ISO. Rats were placed back in the HC and 3 min later tested on the reversal learning session. Groups were the same as the acquisition probe (VEH-VEH, VEH-ISO, PRO-VEH, PRO-ISO). Therefore, if a rat was in a specific group during the acquisition probe it remained in that group for RV1. One hour later, rats in the VEH-VEH group were split in half and were either returned to their HC (VEH-VEH) or given an infusion of ISO (to form the new group VEH-VEH-ISO) and then returned to their HC. This was to assess whether any effect of ISO was due to an enhancement of consolidation. The remainder of the reversal training trials occurred in the absence of any infusions.

### Reversal Probe

Following reversal training, a reversal probe test was given to assess memory for the new escape hole location. The procedure was the same as the curtain probe but without a curtain. The same measures were recorded.

### Histology

Cannula placements were confirmed histologically at the end of the experiments. Following termination of behavioral experiments rats were transcardially perfused with cold 0.1M phosphate buffer solution (PBS) and subsequently cold 4% paraformaldehyde (PFA) in 0.1M PBS. Brains were left to post-fix for 1 hour and then extracted and placed in PFA overnight. The following day they were transferred to a 30% sucrose / 0.1M PBS cryoprotectant solution until saturation. They were then frozen and sectioned using a cryostat to produce 50 μm coronal sections. Every third slice was mounted onto gel-coated slides and Nissl-stained with Cresyl Violet. Slides were then cover slipped and cannula placements were verified under a microscope **(Figure S1).**

### Data Collection and Analysis

Statistical analyses were conducted using SPSS (IBM, version 26) and SigmaPlotTM 11.0 (Systat Software, San Jose, CA). For the DNMP task, the dependent measures were latency to obtain reward, number of errors, and the percentage of trials where a correct choice was made. Latencies were collected using a timer, and the experimenter recorded the number of errors, which was later used to calculate the percentage of correct trials. Pilot data were analyzed using one-way analysis of variance (ANOVA) to compare arm separations 1 through 6. Habituation, and pre-training data were analyzed using two-way (GROUP x DAY) repeated measures (RM) ANOVAs. Acquisition data were analyzed using two-way (ARM-SEPARATION x DAY) RM ANOVAs. Test data were analyzed using two-way RM (GROUP x TRIAL) ANOVAs separately for each arm-separation. Washout, and curtain probe data were analyzed using three-way RM (GROUP x ARM-SEPARATION x DAY) ANOVAs. Pairwise comparisons were made when necessary using Tukey’s HSD test.

In quantifying the EPM data we measured distance traveled, mean speed, time spent immobile, line crossings, time spent in each zone of the maze, and the number of entries into the zones. The locomotor measures (distance, speed, line crossings, and immobility) were analyzed as one-way ANOVAs, and the time spent in each zone, as well as the number of entries, were analyzed using two-way (GROUP x ZONE) RM ANOVAs. Pairwise comparisons were made when necessary using Holm-Sidak tests.

For the Barnes maze data, we measured path efficiency, total distance traveled, latency to reach the escape hole, the number of hole deviations, reference errors, and characterization of the search path used to find the escape hole. Using a Kruskal-Wallis test these data were compared across days during acquisition training. Difference scores in these measures were calculated between the acquisition probe and the last day of acquisition training. For each measure, one-way ANOVAs were then conducted to measure group differences. During each of the probe sessions the maze was divided into 20 equal zones and the time spent in each zone was recorded. Group differences in the time spent in the escape zone, and the escape quadrant (which included the escape zone as well as the two zones to the left and right of the escape zone) were compared using a one-way ANOVA. To demonstrate whether differences in performance existed between the acquisition probe and the curtain probe a two-way (GROUP X SESSION) RM ANOVA was conducted. On the first day of reversal training group comparisons in latency to reach the escape hole, hole deviations, and reference errors were calculated using a one-way ANOVA. During the reversal probe, behavior across groups was analyzed with a one-way ANOVA. Pairwise comparisons were made when necessary using Tukey’s HSD test. In all cases, p < 0.05 was accepted as significant. Error bars in graphs represent +/- sem; *p < 0.05.

## RESULTS

### DNMP pilot experiment: How arm separations were determined

During the pilot experiment rats were habituated to the maze and then taught to obtain a reward in one arm of the maze in under 2 min (sample trials). Once this behavior was acquired, they were given acquisition trials where they were required to remember the location of the previous arm that they had received that reward in (sample trial) and upon the presentation of two open arms, choose the arm they had not entered yet (choice trial) to successfully complete the task and receive an additional reward. Pilot animals were tested on choice trials where the arm separation ranged from 1-6. Across time, animals learned to obtain the reward more quickly (data not shown).

On the last 3 d of training, there were no significant differences in the latency to obtain the reward across all arm separations (F_5,66_ = 1.652, p = 0.159) **(Figure S2A).** However, there were differences in the number of errors made (F_5,66_ = 3.498, p = 0.007) and the percentage of correct trials (F_5,66_ = 3.178, p = 0.012) **(Figures S2B-C)**. Rats made more errors when the arm-separation was 2 arms compared to when the separation was 5 or 6 arms (2 *vs.* 5: p = 0.006; 2 *vs.* 6: 0.026) **(Figure S2B).** When the arm-separation was 2 arms, rats also had the lowest percentage of correct trials, which differed significantly from the 5-arm separation group (p = 0.013) **(Figure S2C)** suggesting that the task was easiest when the separation was 5 arms and most-difficult when the separation was 2 arms. Given that Clelland et al. (2009) reported chance level performance when a 1 arm-separation (45 degrees) was used, and the fact that clockwise and counter clockwise permutations cannot be counterbalanced for a 6-arm separation (180 degrees) in a 12-arm radial maze we decided to use the 2-arm separation for the difficult, DG-dependent, high similarity condition (S2, 60 degrees) and the 5-arm separation for the easier, DG-independent, low similarity condition (S5, 150 degrees). Moreover, these arm-separations were comparable to those used in Clelland et al., (2009) in terms of angular distance.

### The DG orthogonalizes contextual representations when they are highly similar

In the current study, we sought to assess the role of BAR activity during a spatial memory task that relied on the DG. It has been determined (Clelland et al., 2009), that spatial discrimination in an eight-arm radial maze is dependent on the DG when stimuli are presented with little separation but not when presented more widely apart. We adapted this task for the twelve-arm radial maze. Equating angular distance to obtain comparable behavioral results, we concluded optimal arm separations were two- and five arms apart since we saw the greatest difference in latency **(Figure S2A),** number of errors **(Figure S2B),** and percent of correctly conducted trials **(Figure S2C),** demonstrating the S2 condition was more difficult and DG-dependent than the S5 condition. We then assessed the role of BAR activation and inactivation in the DG with ISO and PRO respectively across both S2 and S5 conditions. During habituation, rewards collected increased across days (F_3,114_ = 9.421, p = 0.001) (**Figure S2D)** with significantly more rewards collected on days 3 (p = 0.007) and 4 (p = 0.001) compared to day 1. During pretraining, only sample trials were presented. Animals initially took approximately 200 s to obtain the reward but by the 10^th^ day they could do this in under 30 s (F_9,279_ = 17.45, p = 0.001) with significantly shorter latencies emerging by pre-training day 3 (p = 0.001) **(Figure S2E).** No group differences were observed.

During DNMP acquisition training, animals received both sample and choice trials. By the 6th day, 62.5% of the rats reached criterion; the other 37.5% reached this criterion by day 7. The analysis included the first 3 days and the last 3 days of training data; therefore, the data is inclusive for rats that took 6 days to reach criterion, and for rats that took 7 days, the set excludes the data from day 4. Animals completed the trials more quickly across time (F_5,195_ = 9.041, p = 0.001) and animals in the S2 condition took longer to complete the trials compared to the s5 condition (F_1,195_ = 19.38, p = 0.001) **(Figure S2F).** At the start of acquisition training all rats were performing at an error rate of approximately 50% **(Figure S2G).** As they learned the task, a difference in performance emerged across groups (significant interaction: F_5,195_ = 4.372, p = 0.001) by acquisition day 5 (p = 0.024) with rats improving in the S5 condition to 94% correct trials on day 6 (p = 0.001) whereas this remained low in the S2 condition **(Figure S2G).** A similar pattern emerged for the number of errors made in the choice trials (significant interaction: F_5,159_ = 2.491, p = 0.033) starting on day 5 (p = 0.008) and extending to day 6 (p = 0.001) **(Figure S2H).**

### Biasing memory towards encoding during retrieval can be maladaptive: pre-choice infusions of isoproterenol impaired choice accuracy and increased latency to obtain a reward in a working memory task

Following training, animals were tested on 4 conditions across 4 different testing days **(Figure 1A**, **Table S6).** They were assigned to either the ISO or PRO drug group which, remained consistent across all test days. Differing on each test day was the timing of the infusion: Pre-Sample (PS) or Pre-Choice (PC) **(Figure 1B),** and the arm separation: 2 arms (S2) or 5 arms (S5) **(Figure 1C).** Animals were given a habituation trial (HAB), a baseline trial (BASE; vehicle / saline), and a test trial (TEST, drug). Based on our hypothesis that NE biases the memory system towards encoding, we expected ISO given pre-sample would improve memory while PRO would have the opposite effect, in other words, biasing the system towards encoding when it is adaptive, when learning is occurring. While we found no effect on latency or the number of errors during testing **(Figure 1D & 1F)**, we did find that infusions of PRO resulted in fewer trials where animals made the correct choice (F_1,38_ = 4.897, p = 0.033) potentially due to blockade of NE-driven encoding necessary to complete this task successfully **(Figure 1E).**

Our hypothesis also led us to assume that ISO administered after the sample, and before the choice, when the task required recalling a previously formed contextual map rather than recruiting a new map, would result in impaired memory, in other words, biasing the system when it is maladaptive. As expected, animals in the S2 condition took longer than the S5 condition to obtain the reward (significant interaction: F_1,38_ = 8.416, p = 0.006). More specifically, animals in the ISO group demonstrated longer latencies on the test trial compared to baseline (p = 0.001) and compared to PRO animals (p = 0.001) in the S2 condition. In the S5 condition, ISO animals did not exhibit longer latencies during the test (p = 0.077) **(Figure 1G).** One explanation for these results may be that ISO promotes attentional shifts where animals spend more time exploring the maze rather than focused on the task. While animals did make more errors (and fewer correct trials) when given ISO in both S2 and S5 trials **(Figures 1H-I)**, these effects were not statistically significant. However, this effect was more pronounced in the S2 condition demonstrating the DG is sensitive to this disruption.

### Visuo-spatial learning in the DNMP task depends on contextual elements within the environment

Separating test days, animals received 4 washout days to demonstrate drug clearance. Rats performed similarly to acquisition in terms of latency, percent correct trials, and errors. Likewise, both ISO and PRO animals took longer (F_1,124_ = 17.33, p = 0.001) made more errors (F_1,249_ = 16.88, p = 0.001), and completed less correct trials (F_1,249_ = 13.52, p = 0.001) in the S2 condition **(Figure S3A-F)**. Animals used extra-maze cues to complete the DNMP task demonstrated by comparing data from the curtain probe to the previous washout day. Rats took longer to complete the trials (F_3,249_ = 15.17, p = 0.001) **(Figure S3G),** completed fewer correct trials (F_1,124_ = 31.39, p = 0.001) **(Figure S3H),** and made more errors (F_1,124_ = 31.7, p = 0.001) (**Figure S3I),** in both drug conditions on both S2 and S5 trials during the curtain probe suggesting contextual cues in the room guided their visuo-spatial learning.

### Isoproterenol and propranolol did not affect anxiety-like or locomotor behavior

Here, we sought to determine whether the effects of BAR activation and blockade observed in experiment 1 could be attributed to anxiety-like or locomotor behavior by examining the effect of bilateral DG infusions of ISO and PRO using the EPM. We found no group differences in distance traveled **(Figure 2A),** mean speed **(Figure 2B),** number of line crossings **(Figure 2C),** or time spent immobile **(Figure 2D).** Moreover, all animals made more entries into the open arms (F_1,13_ = 14.76, p = 0.002) **(Figure 2E)** and spent more time in the open arms compared to the closed arms or the start area of the maze (F_2,26_ = 28.04, p = 0.001) **(Figure 2F).** These data indicate the increase in latency and number of errors observed in the DNMP task following infusion of ISO was not due to an increase in locomotion. Moreover, the doses of ISO and PRO we chose did not induce any anxiogenic effects.

**Figure 2.**
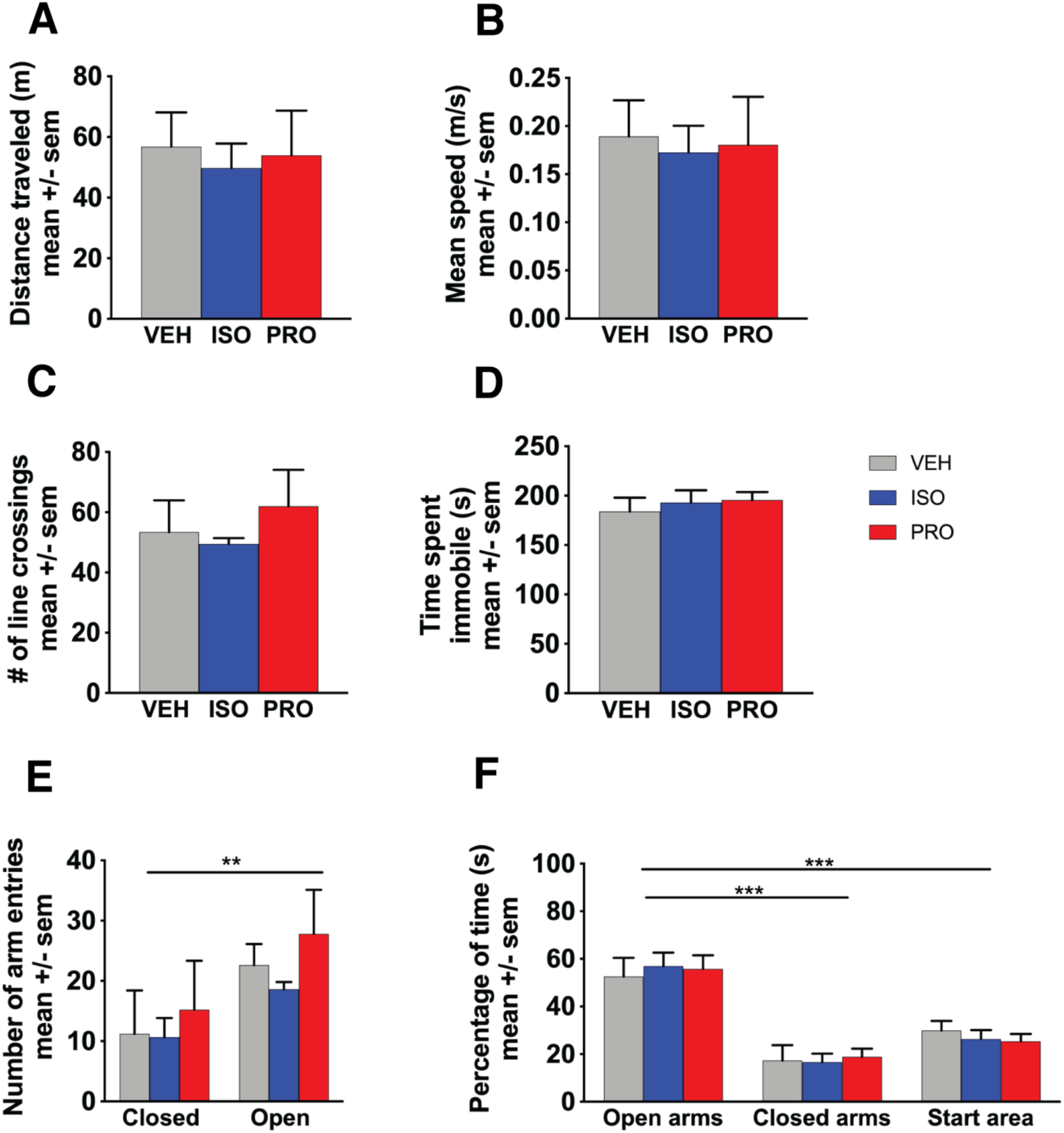
Rats were tested in the EPM to determine if either ISO or PRO affected locomotor behavior. We measured (A) total distance traveled (B) mean speed (C) number of line crossings and (D) time spent immobile and found no effect of either VEH (grey; n = 5), ISO (blue; n = 5) or PRO (red; n = 5). Rats were also tested in the EPM to determine if either ISO or PRO affected anxiety-like behavior. We measured (E) percentage of time the animals spent in the open arms, closed arms, and the start area. We also measured (F) the number of arm entries into the open and closed arms. We found no group differences and no effect of ISO or PRO on anxiety-like behavior. All animals spent more time in the open arms. Significant differences denoted with asterisks * p < 0.05, **p < 0.01, ***p < 0.001. VEH = Vehicle (saline) ISO = Isoproterenol, PRO = Propranolol, EPM = Elevated plus maze.

### Biasing memory towards encoding during retrieval can be maladaptive: infusions of isoproterenol impaired choice accuracy and increased latency to obtain a reward in a reference memory task

Next, we trained animals on a spatial reference memory task using the Barnes maze **(Figure 3A-B).** During this task, rats were placed on a brightly lit circular maze with 20 equidistant holes around the perimeter, one of which led to a dark escape box. The natural tendency for rodents to seek out dark, closed spaces motivated them to escape the open platform into the recessed escape box. The location of the escape hole remained constant. There was a significant decrease in the latency it took to reach the escape hole by the ninth day compared to the first day (χ^2^ = 60.235, p = 0.001, df = 11) **(Figure 3C).** By day 7, animals made fewer hole deviations (χ^2^ = 62.885, p = 0.001, df = 11) **(Figure 3D)** and reference errors (χ^2^ = 73.9, p = 0.001, df = 11) **(Figure 3E).** They also started using a spatial search strategy rather than a random or serial search strategy (χ^2^ = 54.152, p = 0.001, df = 11) **(Figure 3F)** as well as a more efficient path as they exhibited a more direct heading towards the escape hole (χ^2^ = 53.46, p = 0.001, df = 11) **(Figure 3G).** This was further demonstrated by the decrease in total distance traversed per trial by day 9 (χ^2^ = 55.716, p = 0.001, df = 11) **(Figure 3H).**

We next assessed the effect of BAR activation and blockade on spatial reference memory in the acquisition probe test. Although our infusions were aimed at the DG, to demonstrate specificity for, and control for any off-target effects, we included a control group that received both ISO and PRO in addition to our VEH group. To minimize competitive binding, we infused PRO 15 min before the test and ISO 7 min before the test. We compared the last day of acquisition training to performance during the acquisition probe across groups using a difference score for latency **(Figure 3I),** hole deviations **(Figure 3J),** search strategy **(Figure 3K),** and reference errors **(Figure 3L)**. All animals received 2 infusions, either VEH or PRO, 15 min before the probe trial, and then either VEH or ISO, 7 min before the trial. Similar to experiment 1, animals in the VEH-ISO group took longer to reach the escape hole (F_3,13_ = 4.041, p = 0.031) driven by a difference between the VEH-ISO group and the VEH-VEH group (p = 0.031). The VEH-ISO group also had more hole deviations, reference errors, and fewer of these animals used a spatial search strategy, however, these effects did not reach significance. We examined time spent in each of the 20 zones throughout the acquisition probe. Animals in the VEH-ISO group spent greater than chance levels of time in the escape zone (F_3,13_ = 56.74, p = 0.001), but less time compared to other groups (F_3,13_ = 3.128, p = 0.062). These animals also spent less time in the escape quadrant (significant group x chance interaction: F_3,13_ = 3.8, p = 0.037) **(Figure 3M-P)** demonstrating the ISO infusion given 7 min prior to the acquisition probe trial, impaired spatial performance in the maze. This effect was not observed in the other groups.

### Visuo-spatial learning in the Barnes maze depends on contextual elements within the environment

Here, we sought to confirm whether animals used intra or extra-maze cues to locate the escape box. Following 2 days of re-training, we compared the previous trial (re-training trial 2) to the curtain probe and calculated a difference score. The curtain probe was identical to the acquisition probe except animals did not receive drug infusions and a curtain was hung around the maze blocking visual access to contextual cues. As expected, there were no group differences, but all groups showed impaired performance including increased latency (F_1,13_ = 14.85, p = 0.002) **(Figure S4A),** hole deviations (F_1,13_ = 16.92, p = 0.001) **(Figure S4B),** and reference errors (F_1,13_ = 14.03, p = 0.002) **(Figure S4C)**, as well as a decrease in the percentage of animals using a spatial search strategy (F_1,13_ = 21.99, p = 0.001) **(Figure S4D)**. We found rats spent less time in the escape zone (significant interaction: F_3,13_ = 3.869, p = 0.001) and the escape quadrant (Session: F_1,13_ = 40.81, p = 0.001; Group: F_3,13_ = 5.103, p = 0.015) during the curtain probe compared to the acquisition probe, with the exception of rats in the VEH-ISO group, which showed impaired performance on this measure during both tests **(Figure S4E-F).** Time spent in each zone during the curtain probe is shown in **Figure S4G-J**. These results suggest that rats relied on extra-maze cues, rather than interoceptive or intra-maze cues to locate the escape hole.

### Biasing memory towards encoding when new information is present can be adaptive: infusions of isoproterenol improved reversal learning

Following the curtain probe, animals received 2 more days of retraining to reduce possible extinction effects. Next, we assessed the effect of moving the location of the escape hole 180 degrees. To investigate group differences during the first day of reversal learning animals received the same drug treatments as the acquisition probe. We examined latency **(Figure 4A),** reference errors **(Figure 4B)** and hole deviations **(Figure 4C).** Search strategy was not analyzed (all animals used a serial search strategy). We observed no differences in reference errors or hole deviations, however, consistent with our previous results, VEH-ISO animals again, took longer than the other groups to find the new escape box (F_3,13_ = 5.48, p = 0.012). Post-hoc analyses showed that this was due to a difference between VEH-ISO rats and PRO-VEH rats (p = 0.019) as well as VEH-VEH rats (p = 0.021). Following this trial, half the animals in the VEH-VEH group were given an additional infusion of ISO (VEH-VEH-ISO) to rule out potential effects on memory consolidation rather than remapping specific to the time of learning.

**Figure 4.**
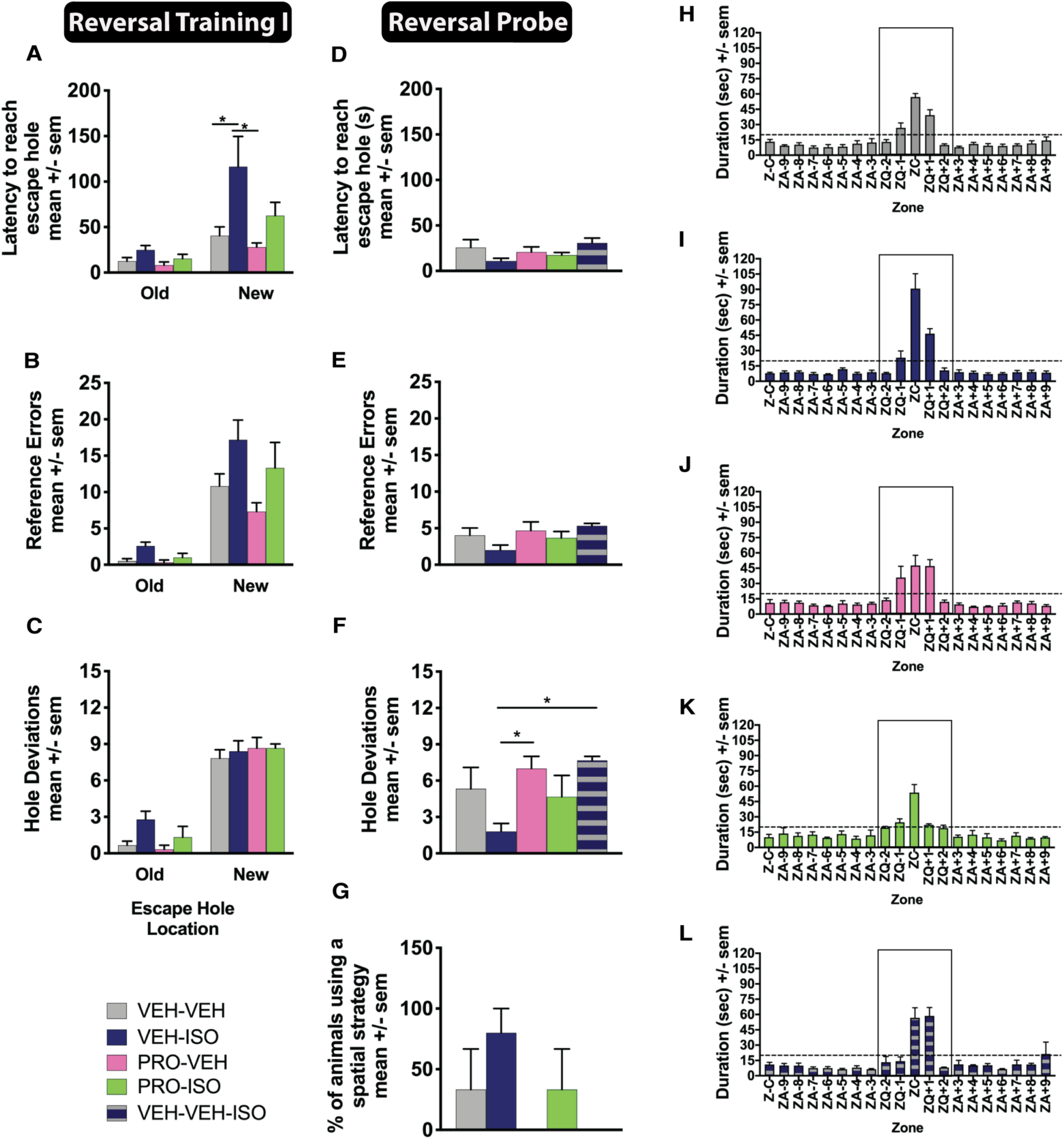
On the first day of Reversal training, we assessed the effect of moving the location of the escape hole 180 degrees. 15 min prior to the first reversal training session, rats were given an infusion of either saline or PRO. Seven min prior to the session rats were given another infusion of either saline or ISO. Groups were the same as the acquisition probe [VEH-VEH = grey (n = 6), VEH-ISO = blue (n = 5), PRO-VEH = pink (n = 3), PRO-ISO = green (n = 3)]. One hour later after the session ended, rats in the VEH-VEH group were split in half and were either returned to their home cage or given an infusion of ISO to form the new group VEH-VEH-ISO [blue with grey stripes (n=3)] and then returned to their home cage. This was to assess whether any effect of ISO was due to an enhancement of consolidation. We calculated (A) latency, (B) reference errors, and (C) hole deviations for both the old and new escape holes. After 5 days of reversal training, animals were given a 5-min reversal probe test where the escape box was removed and replaced with a false escape box. We measured (D) latency, (E) reference errors, (F) hole deviations, and (G) percentage of animals using a spatial search strategy. (H-L) During the reversal probe test, the maze was divided into 20 equal zones. The escape zone (ZC) contained the escape hole, and the escape quadrant (ZC, ZQ-2, ZQ-1, ZQ+1, ZQ+2) contained the escape zone plus the 2 zones to the left and right of the escape zone. Time spent in the escape zone and quadrant were calculated. Chance levels are expressed as a dotted line. Significant differences are denoted with asterisks * p < 0.05, **p < 0.01, ***p < 0.001. ISO = Isoproterenol, PRO = Propranolol, VEH = Vehicle (saline), ZC = Escape zone, ZQ = Escape quadrant.

For the remainder of the reversal training days, rats received 2 trials / day. We computed the mean and analyzed learning for the new location of the escape box by comparing performance across days. We looked at latency, hole deviations, search strategy, and reference errors. Similar to acquisition training, we also looked at total path efficiency and distance traveled. For all analyses, we found no group differences but did find across days animals learned the new escape box location (Latency: F_3,36_ = 6.669, p = 0.001; Hole Deviations: F_3,36_ = 11.95, p = 0.001; Reference Errors: F_3,36_ = 13.64, p = 0.001; Spatial Strategy: F_3,36_ = 5.533, p = 0.003; Path Efficiency: F_3,36_ = 7.731, p = 0.001; Distance Traveled: F_3,36_ = 10.1, p = 0.001) (data not shown).

To assess memory for the new escape hole location, as well as investigate whether activation of BARs in the DG prior to reversal training conferred any mnemonic advantage, we examined the same dependent measures in the reversal probe test. There were no group differences in latency **(Figure 4D).** Animals in the VEH-ISO group had fewer hole deviations (F_4,12_ = 4.828, p = 0.015) compared to the PRO-VEH (p = 0.035) and VEH-VEH-ISO group (p = 0.017) **(Figure 4F)** suggesting their spatial map for finding the new escape hole was more refined compared to animals that had not received ISO prior to reversal learning. Moreover, we confirmed that the beneficial effect conferred via ISO was not due to an enhancement of memory consolidation. No group differences were observed in terms of reference errors made **(Figure 4E)** or the search strategy used **(Figure 4F).** The distribution of time spent in each of the 20 zones during the reversal probe is shown in **Figure 4H-L.** Animals spent greater than chance levels in the escape zone (F_1,13_ = 69.93, p = 0.001), however, there was no group differences. With respect to time spent in the escape quadrant, all rats spent greater than chance levels in the escape quadrant (F_1,13_ = 263.7, p = 0.001), however, VEH-ISO animals showed enhanced performance as they spent more time in the escape quadrant compared to other groups. This effect did not reach significance (F_3,13_ = 3.319, p = 0.054), but may reflect a level of enhanced cognitive flexibility imparted via the activation of BARs in the HPC, prior to learning a new escape hole location. Post-hoc analyses revealed a significant difference between the VEH-ISO group and the VEH-VEH group (p = 0.007) and the PRO-ISO group (p = 0.003). Importantly, drug treatments did not have any effect on locomotor behavior as we compared the total distance traversed in the maze during each of the probes and found no group differences **(Figure S5A-C).**

## DISCUSSION

We previously demonstrated that novelty-associated LC activation causes the HPC to recruit a new population of neurons to represent contextual change. Based on these results as well as the established role of the LC in processing novelty (Berridge & Waterhouse, 2003; Harley 2007), we hypothesized that NE may serve as a neuromodulatory signal involved in updating memories at the ensemble level. The aim of this study was to show that if NE can shift the HPC towards encoding, this could be adaptive if an animal is presented with new information and requires contextual representations to be updated (e.g., reversal learning), but this could also be maladaptive, if the animal requires reactivation of a previously formed contextual map to complete a task (e.g., reference and working memory). We sought to test this by investigating the involvement of the noradrenergic system, and more specifically BARs, in spatial working memory using the DNMP task, and spatial reference memory and reversal learning using the Barnes maze. We found intra-DG infusions of ISO given prior to retrieval (when a contextual map formed during encoding needs to be reactivated) caused deficits in working and reference memory retrieval, an effect blocked by pre-treatment with PRO. In contrast, we also found intra-DG infusions of ISO given prior to reversal learning, conferred a slight advantage to the animal improving performance during the probe test. These results complement our previous findings showing phasic LC activation, and subsequent NE release associated with the detection of novelty, induces HPC plasticity involving the recruitment of new neurons and a global reorganization of contextual representations (Bouret and Sara, 2005; Grella et al., 2019). Together, these data suggest the presence of NE acts as a molecular switch to bias memory processing in the direction of encoding, while its absence promotes retrieval.

Consistent with previous work (Clelland et al., 2009), animals performed better when the distance between arms in the radial maze was greater. Curtain probe trials confirmed animals relied on extra-maze cues to complete the task, cues which tend to be more disparate when the distance between arms is greater. In contrast, when the arms are closer, distal cues used to orient potentially overlap causing interference in the representations for the arms requiring the DG to orthogonalize these representations. As expected, pre-choice administration of ISO resulted in impairments via increased latency and decreased choice accuracy. These effects were more pronounced in the S2 condition compared to S5, consistent with the role of the DG in pattern separation. It is possible HPC-dependent recollection was impaired and performance here relied solely on HPC-independent familiarity mechanisms (Eichenbaum et al., 2012).

Throughout the study, ISO increased latency to the goal location which, presumably involved enhanced motivation to explore the environment, a hypothesis derived from previous studies (Flicker and Geyer, 1982; Geyer and Masten, 1989), where an increase in *diversive* (reconnaissance-like) exploration developed following bilateral infusions of ISO in the DG. The characteristics of DG neuronal ensemble activity during exploration currently remain unclear. Importantly, increased latency was not due to changes in locomotion. At a similar dose, ISO has been shown to mediate anti-depressant effects (Zhang, Frith, and Wilkins, 2001), however, it is not likely this would explain the increases in latency. Similarly, these effects were not the result of changes in anxiety-related behavior given all animals spent more time in the open arms of the EPM, suggesting both ISO and PRO do not impart anxiogenic effects, an important determination since altered BAR function and heightened noradrenergic tone have been linked to anxiety in the past (Leo, Guescini, Genedani, Stocchi, Carone, Filaferro et al., 2015). In contrast, PRO is associated with attenuated physiological responses to anxiety (Nandhra, Murphy and Sule, 2013; Argolo, Cavalcanti-Ribeiro, and Netto, 2015).

The literature regarding the role of NE on memory is heavily focused on consolidation and reconsolidation of emotional memory (Cahill et al., 1994; van Stegeren et al., 1998; Przybyslawski et al., 1999). In humans, activation of BARs results in augmented consolidation which is thought to be mediated via receptors in the amygdala (Cahill et al., 1994; Cahill and McGaugh, 1998; van Stegeren et al., 1998; Soeter and Kindt, 2012; Chamberlain and Robbins, 2013; Barsegyan et al., 2014; Kuffel et al., 2014 for review see Ferry et al., 1999; McGaugh, 2000; Roozendaal et al., 2009; Roozendaal and McGaugh, 2011), an effect which disappears if participants are pretreated with PRO (Cahill et al., 1994; van Stegeren et al., 1998; Maheu et al., 2004) or if given PRO post-learning (Sara et al., 1999; Tronel et al., 2004; Roozendaal et al., 2008; Barsegyan et al., 2014). Fewer studies have examined the role of NE on mechanisms of encoding and retrieval (Brown and Silva, 2004; Chamberlain et al., 2006; Thomas, 2015; Rimmele et al., 2016). Although NE is generally thought to enhance encoding and retrieval, it is difficult to differentiate whether activation of the NE system is affecting encoding or consolidation per se in situations where emotional memories are remembered better than neutral memories. Furthermore, these effects are also amygdala-dependent and not necessarily mediated by the HPC.

Here we report that rats were impaired on memory retrieval following activation of BARs in the DG, when tested following drug administration, consistent with our hypothesis. We also report blockade of BARs had no effect. This finding was expected since the DNMP choice phase required reactivation of a previously formed representation and did not require recruitment of a new ensemble, therefore, blocking the recruitment of new neurons via BARs at this point in the task would theoretically have no effect on retrieval. This result is consistent with a previous study (Qi et al., 2008) that demonstrated that even with a much larger dose (15μg), PRO had no effect on contextual fear memory retrieval after 1- or 7 d following learning. Moreover, they found ISO (10μg) infused 30 min before the retention test disrupted retrieval of 7-day contextual fear memory, consistent with our findings. In rats, PRO has led to deficits in spatial reference memory in the water maze (Qi et al., 2008), disrupted the retrieval of a cocaine-associated memory (Otis and Mueller, 2011), and abolished cocaine conditioned place preference (Fitzgerald et al., 2016). However, in humans, PRO given prior to a test of memory retrieval had no effect (Rimmele et al., 2016). When our manipulations were conducted prior to encoding rather than retrieval, our effects overall were less pronounced, however, we did see a slight impairment with PRO consistent with previous work (Rimmele et al., 2016). One important difference between these studies and our DNMP experiment is we did not look at memory retrieval post-consolidation but instead examined the role of BARs on working memory.

The literature with respect to working memory is less consistent. In rodents, LC inactivation (Khakpour-Taleghani et al., 2009) did not affect spatial working memory. Administration of PRO in rats also had no effect (Kobayashi et al., 1995; Ohno et al., 1997). In Rhesus monkeys, moderate doses of PRO (0.01, 0.05 and 0.1 mg/kg) impaired spatial working memory, while a low dose (0.005 mg/kg) and high dose (0.5 mg/kg) had no effect (Wang et al., 2012). In humans, a low dose (25 mg) of PRO impaired numerical working memory (Müller et al., 2005), and repeated administration of a high (160 mg) dose impaired working memory (Frcka and Lader, 1988). Several other studies using a moderate dose (40 mg) found no effect at all of PRO on working memory in humans (Bodner et al., 2012; Becker et al., 2013; Ernst et al., 2016). Therefore, the role of NE on working memory is still unclear.

Our investigation was focused on BARs given that earlier work demonstrated that novelty-associated phasic LC activation resulted in BAR-dependent downstream DG plasticity (Pickel et al., 1974; Kitchigina et al., 1997). ISO is a non-selective BAR agonist, which has almost no affinity to α-adrenergic receptors (Copik, Baldys, Nguygen, Sahdeo, Ho, Kosaka et al., 2015) therefore the effects we observed are not likely attributed to activation of this receptor subtype. In contrast, PRO is a non-selective BAR antagonist, with weak inhibitory effects at the NE transporter causing NE to pool in the synapse, which can result in activation of α-adrenergic receptors (Tuross and Patrick, 1986). However, given the lack of effect in the PRO group, it is unlikely α-adrenergic receptors played a role here.

When we tested reference memory in the Barnes maze, administration of ISO prior to the acquisition probe resulted in longer latencies to reach the escape hole, and less time spent in the escape quadrant and zone. Animals in the ISO group made more errors, and the number of hole deviations was greater, however, these measures were not significant. Effects were blocked by PRO. During the curtain probe, performance was impaired across groups on all measures confirming animals used extra-maze cues to orient. We compared performance across groups during the acquisition probe and the curtain probe to show the level of impairment induced following administration of ISO during the acquisition probe, was equal in magnitude to the impairment induced if there were no extra-maze cues present to successfully perform the task. In both cases, the animal’s “map” needed to solve the task was compromised. These results are consistent with the DNMP data suggesting BAR activation in the DG immediately prior to a test of memory retrieval impairs memory.

When animals were infused with ISO a second time, prior to reversal training, they again exhibited longer latencies to reach the new escape hole. However, these same animals showed enhanced performance in terms of number of hole deviations, and time spent in the new escape zone compared to other groups during the reversal probe. These effects were blocked by PRO and were not due to differences in locomotor activity. This is consistent with previous research (Segal and Edelson, 1978) and the hypothesis that the modulating effect of NE depends on the stage of training where NE promotes cognitive flexibility. The noradrenergic system has been previously implicated in reversal learning (Aston-Jones et al., 1994, 1997; Rajkowski et al., 1994; Aston-Jones and Cohen, 2005) and in shifting attention to environmental imperatives (Sara, 2009). Here we see that activation of BARs can confer a slight advantage to the animal promoting a disengagement from established representations and the recruitment of new representations towards an enhancement of processes that promote the incorporation of new information (Bouret and Sara, 2005; Harley, 2007).

The current work highlights the involvement of hippocampal BARs in determining the flexibility of contextual representations to promote new learning in a way that supports adaptive behavior. These data suggest targets for anxiety disorders such as PTSD, which are characterized by noradrenergic dysregulation (Hendrickson and Raskind, 2016), and may also involve impairments in memory updating mechanisms where the incorporation of new information is not effectively encoded. This encoding failure may represent an inability to remap (i.e., reactivating trauma-related representations rather than incorporating safety signals into existing memories through remapping processes) (Maren et al., 2013; Morrison and Ressler, 2014; Giustino et al., 2016; Liberzon and Abelson, 2016; Elsey and Kindt, 2017; Lee et al., 2017; Sheynin and Liberzon, 2017). Therefore, further understanding of the neurobiological mechanisms involved in updating memories may provide insight into novel treatment strategies.

## ACKNOWLEDGEMENTS

This work was supported by NSERC grants awarded to SLG and DFM, and an OMHF grant awarded to DFM.

## SUPPLEMENTAL FIGURES & FIGURE LEGENDS

**Figure S1.**
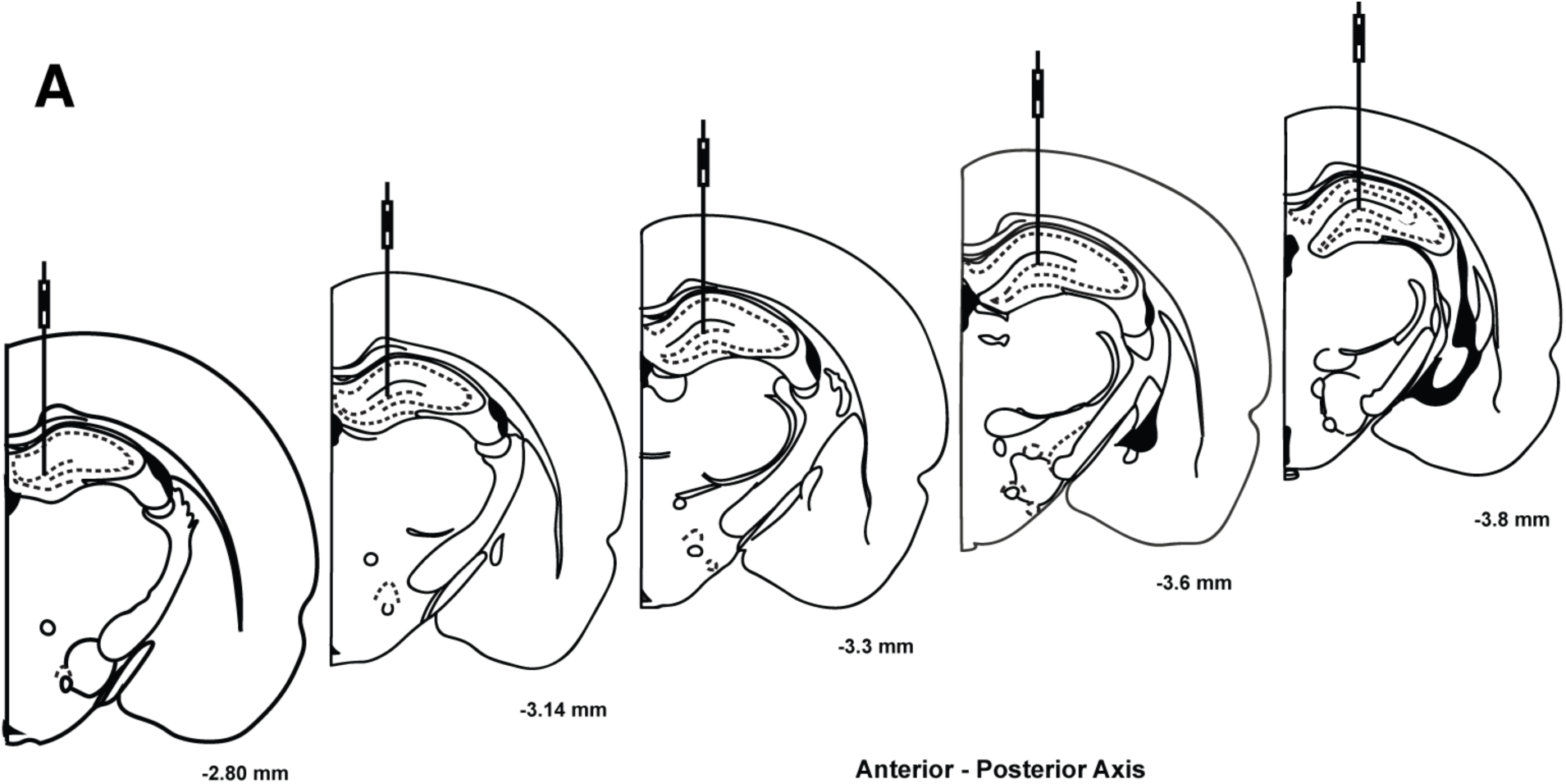
Following termination of behavioral experiments rats were transcardially perfused and brains were extracted, sectioned, and slices were mounted onto slides and Nissl-stained with Cresyl Violet so that cannula placements could be verified histologically under a microscope. For all rats included in the analyses of all experiments, cannula tracts were placed between -2.8 and -3.8mm along the anterior-posterior (AP) axis relative to Bregma using the Paxinos and Watson (2013) rat brain atlas.

**Figure S2.**
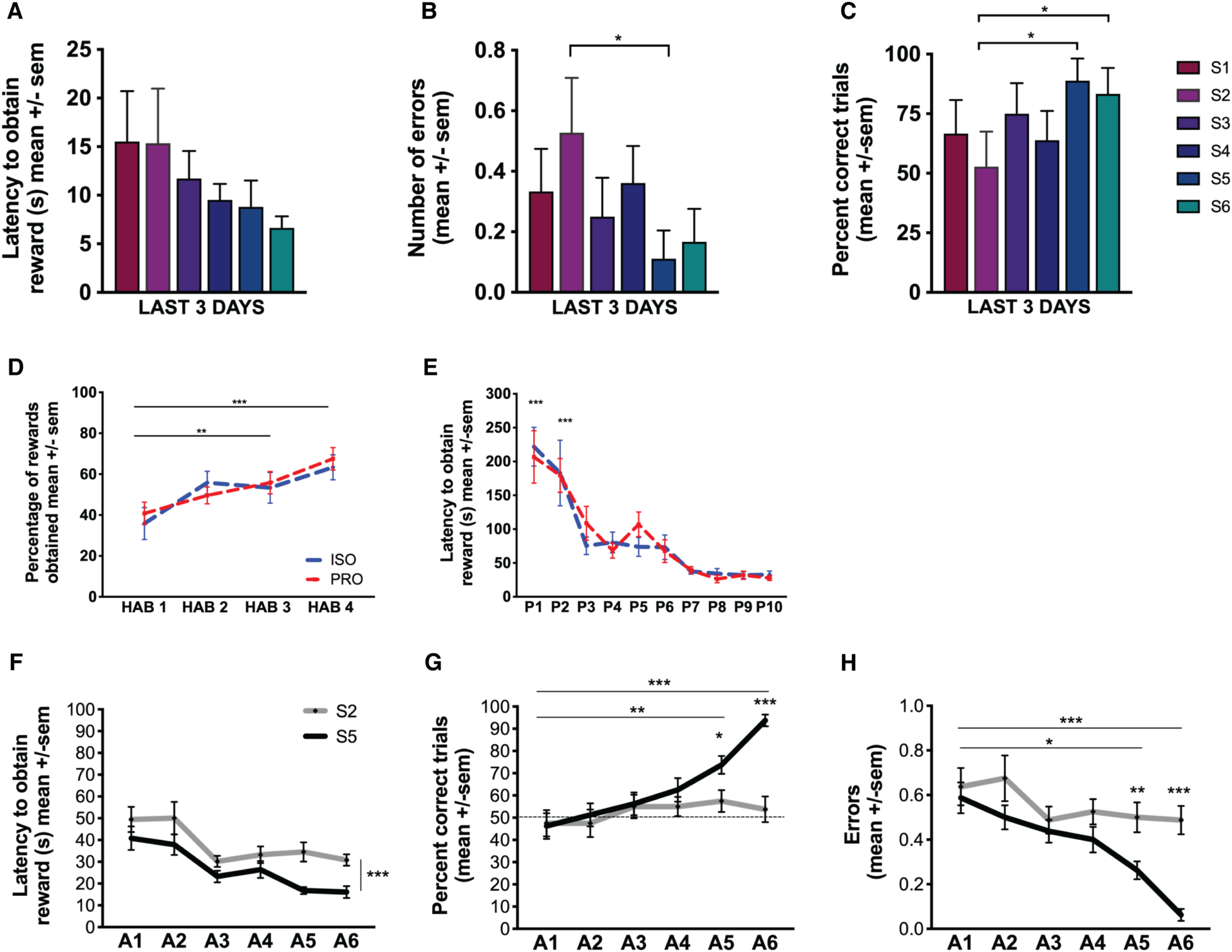
We ran a pilot to determine which arm separations would yield the most comparable results, in terms of behavior and angular distance, to the study from which the DNMP task was adapted. Animals (n = 12) received habituation and pre-training trials (not shown) prior to acquisition-training to assess performance on arm separations 1-6 (S1-6). Dependent measures were: (A) latency to obtain the reward (B) percentage of correct trials and (C) number of errors made. Data is shown for the last 3 days. The most disparate performance was between the 5-arm (S5, 150 degrees, DG-independent, royal blue) and 2-arm conditions (S2, 60 degrees, DG-dependent, magenta). Experiment 1: Prior to acquisition training, animals received 4 days of (D) habituation (HAB 1-4) and then 10 days of (E) pre-training (P1-10) where they were trained to obtain a reward from the maze in under 2 min. (F-H) Acquisition training (A1-A6) lasted 6 days and by the 5^th^ day a difference emerged where rats performed better when they were tested in the S5 condition (black) compared to the S2 condition (grey) (n = 17) demonstrating that the S2 condition was more difficult. Significant differences are denoted with asterisks * p < 0.05, **p < 0.01, ***p < 0.001.

**Figure S3.**
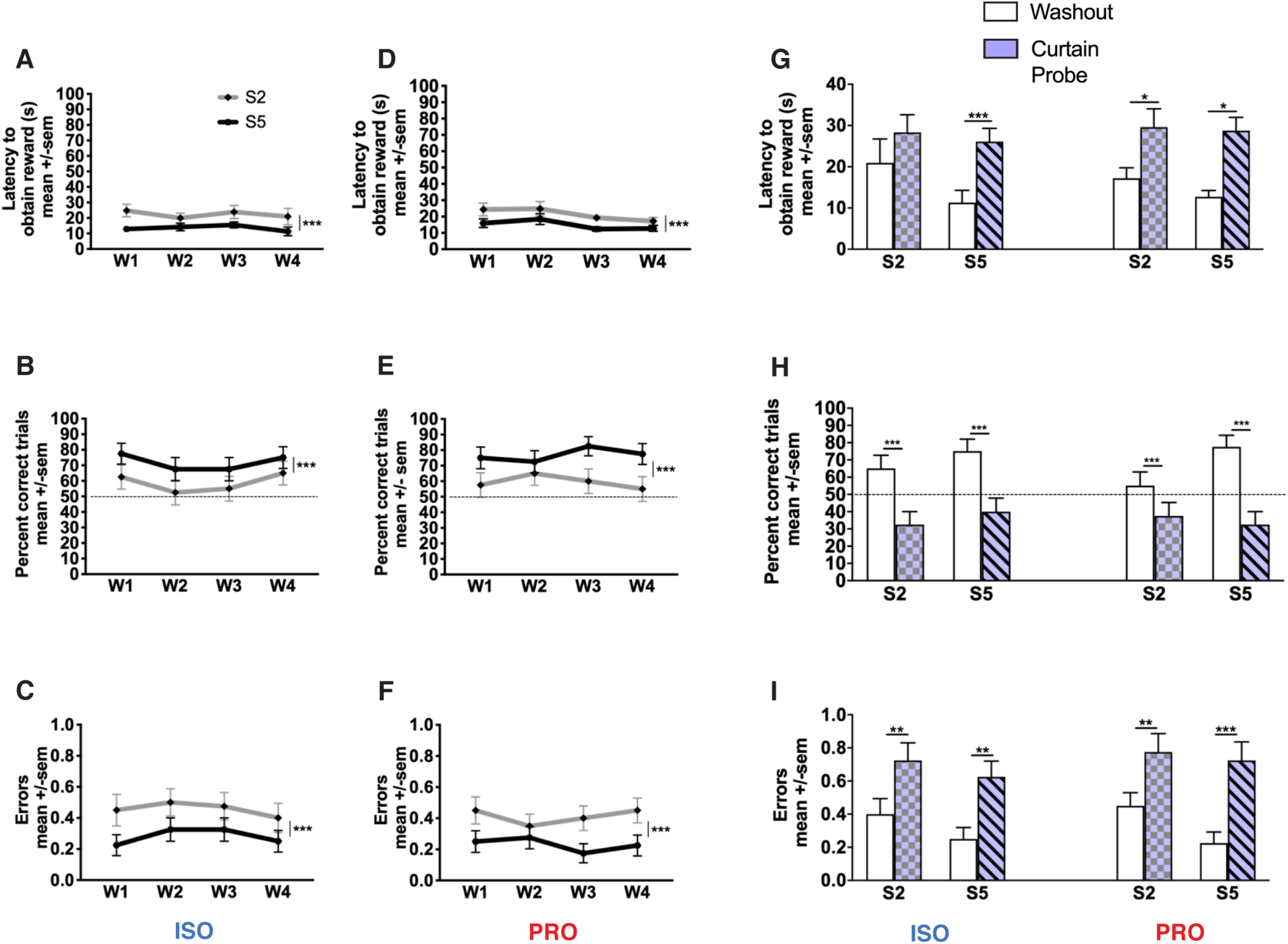
(A-F) Separating the test days, animals tested on S2 (grey; n = 17) and S5 (black; n = 17) conditions were given 4 washout days to ensure the drug had cleared. Dependent measures during washout sessions were latency to obtain reward, percent correct trials and number of errors. Rats were given either isoproterenol (left) or propranolol (right). (G-I) A curtain probe was administered to ensure rats were using extra-maze cues. Compared to the washout (white), during the curtain probe (purple) rats took longer to obtain the reward, demonstrated fewer percent correct trials and made more errors. Significant differences are denoted with asterisks * p < 0.05, **p < 0.01, ***p < 0.001, W1-W4 = Washout sessions 1-4.

**Figure S4.**
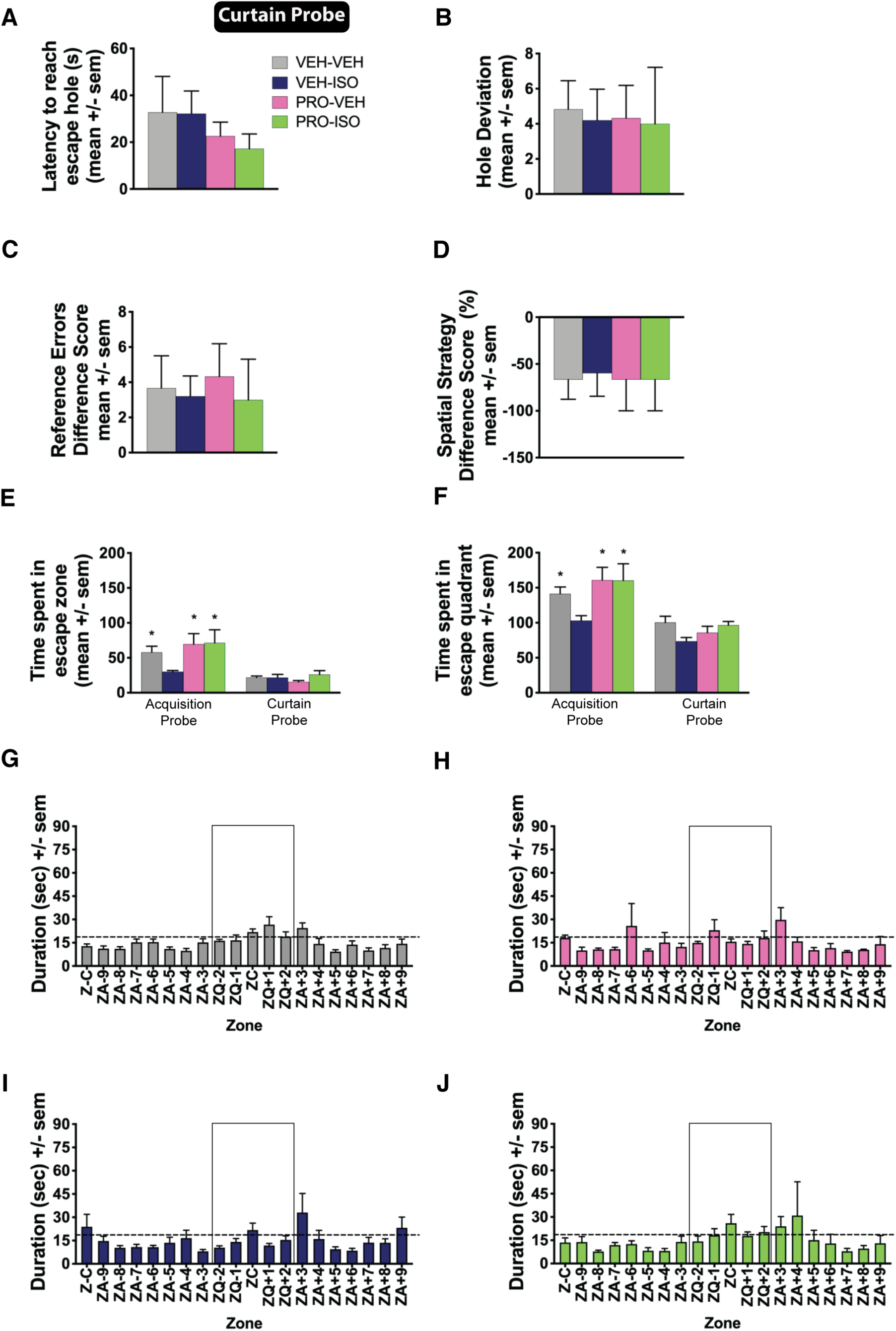
Animals were given a curtain probe trial to ensure they were using extra-maze cues to locate the escape hole. We compared performance during the last trial prior to this (re-training II) to the curtain probe and calculated a difference score. All animals showed impaired performance including increased (A) latency, (B) hole deviations, and (C) reference errors. (D) The percentage of animals using a spatial search strategy also decreased demonstrating that they were using extra-maze cues to solve the task. During the curtain probe, the maze was divided into 20 equal zones. The escape zone contained the escape hole (ZC), and the escape quadrant contained the escape zone plus the 2 zones to the left and right of the escape zone (ZC, ZQ-2, ZQ-1, ZQ+1, ZQ+2). Time spent in the (E) escape zone and the (F) escape quadrant was compared across groups [VEH-VEH n = 6 (grey); VEH-ISO n = 5 (blue); PRO-VEH n = 3 (pink); PRO-ISO n = 3 (green)] during the acquisition probe and the curtain probe to show that the level of impairment induced following administration of ISO (VEH-ISO) during the acquisition probe, was equal in magnitude to the impairment induced if there were no extra-maze cues present to successfully perform the task. In both cases, the animal’s “map” needed to solve the task was compromised. (G-J) Time spent in each zone during the curtain probe. Animals were equally impaired across groups and spent an equal amount of time in all zones of the maze. Time spent in the escape zone or quadrant was not greater than chance (dotted line) suggesting that animals used extra-maze cues to locate the escape hole. Significant differences are denoted with asterisks * p < 0.05, **p < 0.01, ***p < 0.001. VEH = Vehicle, ISO = Isoproterenol, PRO = Propranolol, ZC = escape zone, ZQ = escape quadrant.

**Figure S5.**
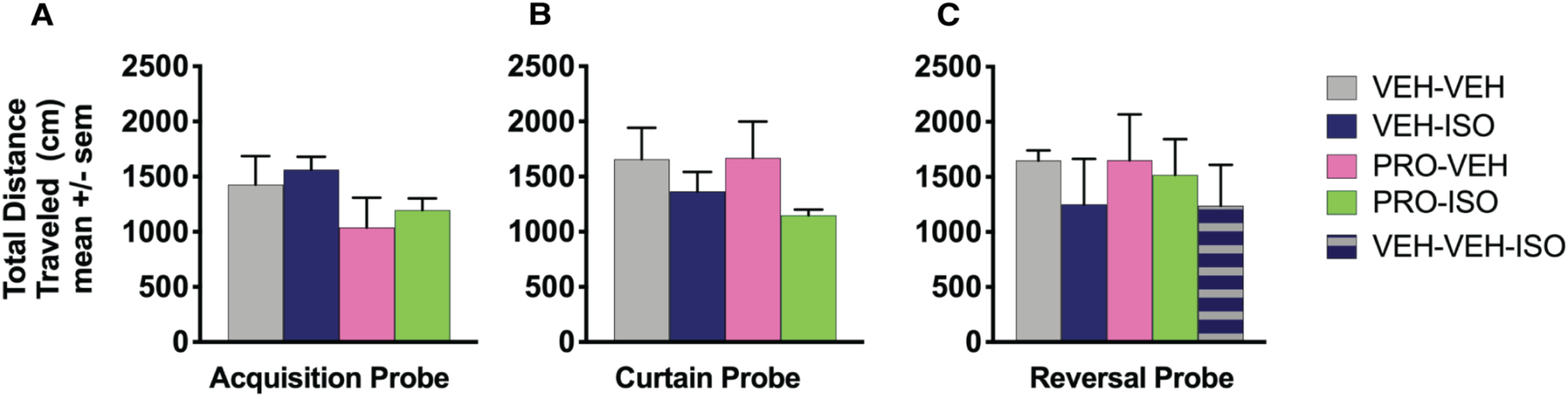
Total distance was measured in each of the probes (A) acquisition probe (B) curtain probe and the (M) reversal probe. Groups: [VEH-VEH n = 6 (grey); VEH-ISO n = 5 (blue); PRO-VEH n = 3 (pink); PRO-ISO n = 3 (green); VEH-VEH-ISO n = 3 (blue with grey stripes)]. Significant differences are denoted with asterisks * p < 0.05, **p < 0.01, ***p < 0.001.

**Table S1.** Number of animals in each experiment.

| Exp | Behavioral Paradigm | Group | n | Figure |
| --- | --- | --- | --- | --- |
| Pilot | DNMP | -- | 12 | S1A-S1C |
| 1 | DNMP | ISO | 20 | 1D-1I; S1D-H; S2A-I |
| 1 | DNMP | PRO | 20 | 1D-1I; S1D-H; S2A-I |
| 2 | EPM | VEH | 5 | 2A-2F |
| 2 | EPM | ISO | 5 | 2A-2F |
| 2 | EPM | PRO | 5 | 2A-2F |
| 3 | BARNES | VEH-VEH | 6 | 3C-3J; 4A-C; S3A-J; S5A-B |
| 3 | BARNES | VEH-ISO | 5 | 3C-3J; 4A-C; S3A-J; S5A-B |
| 3 | BARNES | PRO-VEH | 3 | 3C-3J; 4A-C; S3A-J; S5A-B |
| 3 | BARNES | PRO-ISO | 3 | 3C-3J; 4A-C; S3A-J; S5A-B |
| 3 | BARNES | VEH-VEH-ISO | 3 | 4D-3L; S5C |

Note: DNMP = Delayed Non-Match to Position; EPM = Elevated Plus Maze; BARNES = Barnes Maze; ISO = Isoproterenol; PRO = Propranolol; VEH = Vehicle (Saline).

**Table S2.** List of studies used to determine drug doses

| Reference | Drug | Concentration (mM) | Concentration (µg/µL) | Volume Infused (per side/µL) | Infusion Rate (µL/min) | Duration (min) | Total Mass (per side/µg) | Target | Effects |
| --- | --- | --- | --- | --- | --- | --- | --- | --- | --- |
| Geyer & Masten, 1989 | L-isoproterenol hydrochloride | 0.4000 | 0.1000 | 20.0 | 0.3330 | 60.0 | 2.0000 | DG | increased exploration |
| Geyer & Masten, 1989 | L-isoproterenol hydrochloride | 1.2000 | 0.3000 | 20.0 | 0.3330 | 60.0 | 6.0000 | DG | increased exploration |
| Sun et al., 2005 | L-isoproterenol hydrochloride | 400.0000 | 99.088 | 2.0 | 0.0667 | 30.0 | 198.1760 | CA1 | spatial memory impairments |
| Qi et al., 2008 | L-isoproterenol hydrochloride | 40.3682 | 10.0000 | 1.0 | 0.5000 | 2.0 | 10.0000 | CA1 | impaired retrieval |
| Alsene et al., 2011 | L-isoproterenol hydrochloride | 12.1104 | 3.0000 | 0.5 | 0.5000 | 1.0 | 1.5000 | DG & CA1 | no effect |
| Alsene et al., 2011 | L-isoproterenol hydrochloride | 40.3682 | 10.0000 | 0.5 | 0.5000 | 1.0 | 5.0000 | DG & CA1 | no effect |
| Alsene et al., 2011 | L-isoproterenol hydrochloride | 121.1045 | 30.0000 | 0.5 | 0.5000 | 1.0 | 15.0000 | DG & CA1 | no effect |
| Lethbridge, Walling, & Harley, 2014 | L-isoproterenol hydrochloride | 0.0001 | 0.0000247 | 1.0 | 0.0800 | 12.0 | 0.0000247 | DG | induced LTD |
| Lethbridge, Walling, & Harley, 2014 | L-isoproterenol hydrochloride | 0.0010 | 0.000247 | 1.0 | 0.0800 | 12.0 | 0.000247 | DG | no effect |
| Lethbridge, Walling, & Harley, 2014 | L-isoproterenol hydrochloride | 0.0100 | 0.00247 | 1.0 | 0.0800 | 12.0 | 0.00247 | DG | no effect |
| Lethbridge, Walling, & Harley, 2014 | L-isoproterenol hydrochloride | 0.1000 | 0.0247 | 1.0 | 0.0800 | 12.0 | 0.0247 | DG | induced LTP |
| Hansen & Manahan-Vaughan, 2015 | L-isoproterenol hydrochloride | 16.1473 | 4.0000 | 5.0 | 1.0000 | 5.0 | 20.0000 | ICV | strengthened LTP |
| Garrido-Zinn et al., 2016 | L-isoproterenol hydrochloride | 80.7363 | 20.0000 | 0.5 | 0.5000 | 1.0 | 10.0000 | BLA | no effect |
| Garrido-Zinn et al., 2016 | L-isoproterenol hydrochloride | 40.3682 | 10.0000 | 1.0 | 1.0000 | 1.0 | 10.0000 | CA1 | impaired retrieval |
| Current Study | isoproterenol-bitartrate | 47.3350 | 10.0000 | 0.5 | 0.5000 | 1.0 | 5.0000 | DG | impaired spatial memory |
| Ji et al., 2003 | propranolol hydrochloride | 16.9027 | 5.0000 | 1.0 | 0.5000 | 2.0 | 5.0000 | CA1 | impaired consolidation |
| Chai et al., 2014 | propranolol hydrochloride | 33.8055 | 10.0000 | 0.5 | 0.5000 | 1.0 | 5.0000 | CA1 | blocked NE-facilitated memory enhancements |
| Hatfield & McGaugh, 1999 | propranolol hydrochloride | 5.0708 | 1.5000 | 0.2 | 0.5000 | 0.4 | 0.3000 | BLA | spatial memory impairments |
| Hansen & Manahan-Vaughan, 2015 | propranolol hydrochloride | 1.3522 | 0.4000 | 5.0 | 1.0000 | 5.0 | 2.0000 | ICV | LTP impairment |
| Straube et al., 2003 | propranolol hydrochloride | 0.0068 | 0.0020 | 5.0 | 1.2500 | 4.0 | 0.009998 | ICV | blocked novelty induced LTP reinforcement in DG |
| Walling & Harley, 2004 | propranolol hydrochloride | 20.2840 | 6.0000 | 5.0 | 1.0000 | 5.0 | 30.0000 | ICV | blocked LC-GLUT induced plasticity |
| Qi et al., 2008 | propranolol hydrochloride | 50.7082 | 15.0000 | 1.0 | 0.5000 | 2.0 | 15.0000 | CA1 | did not impair retrieval |
| Barsegyan, Mcgaugh, & Roozendaal, 2014 | propranolol hydrochloride | 1.6903 | 0.5000 | 0.2 | 0.4000 | 0.5 | 0.1000 | BLA | no effect |
| Barsegyan, Mcgaugh, & Roozendaal, 2014 | propranolol hydrochloride | 5.0710 | 1.5000 | 0.2 | 0.4000 | 0.5 | 0.3000 | BLA | diminished NE-facilitated memory enhancements |
| Current Study | propranolol hydrochloride | 10.1420 | 3.0000 | 0.5 | 0.5000 | 1.0 | 1.5000 | DG | no effect |

Note: DG = Dentate Gyrus; CA1= Cornu Ammonis; BLA = Basolateral Amygdala; ICV = Intracerebroventricular; LC = Locus Coeruleus; NE = Norepinephrine; GLUT = Glutamate; LTP = Long Term Potentiation; LTD = Long Term Depression. All hippocampal regions (DG & CA1) were dorsally targeted. All injections were bilateral except ICV injections.

**Table S3.** Combinations of arm separations used for acquisition and washout.

| 5 CW |  | 5 CCW |  | 2 CW |  | 2 CCW |  | Day |
| --- | --- | --- | --- | --- | --- | --- | --- | --- |
| Sample Arm | Choice Arm | Sample Arm | Choice Arm | Sample Arm | Choice Arm | Sample Arm | Choice Arm |  |
| 2 | 7 | 11 | 6 | 3 | 5 | 12 | 10 | A1 |
| 11 | 4 | 8 | 3 | 7 | 9 | 6 | 4 | A2 |
| 6 | 11 | 9 | 4 | 12 | 2 | 5 | 3 | A3 |
| 7 | 12 | 10 | 5 | 6 | 8 | 11 | 9 | A4 |
| 9 | 2 | 4 | 11 | 8 | 10 | 7 | 5 | A5 |
| 3 | 8 | 12 | 7 | 9 | 11 | 4 | 2 | A6 |
| 12 | 5 | 3 | 10 | 5 | 7 | 2 | 12 | W1 |
| 4 | 9 | 7 | 2 | 10 | 12 | 9 | 7 | W2 |
| 10 | 3 | 5 | 12 | 4 | 6 | 10 | 8 | W3 |
| 5 | 10 | 2 | 9 | 2 | 4 | 8 | 6 | W4 |

Note: Animals were tested with an arm separation of 2 or 5 counterbalanced for clockwise (grey) and counter-clockwise (white) directions. The order of testing is shown in Table S5.

**Table S4.** Testing Schedule: Order of S2 and S5 trials during acquisition and washout.

| Rat | Trial 1 | Trial 2 | Trial 3 | Trial 4 |
| --- | --- | --- | --- | --- |
| 1 | 5R | 2L | 2R | 5L |
| 2 | 5R | 2L | 5L | 2R |
| 3 | 5R | 2R | 2L | 5L |
| 4 | 5R | 2R | 5L | 2L |
| 5 | 5R | 5L | 2R | 2L |
| 6 | 5R | 5L | 2L | 2R |
| 7 | 5L | 2L | 2R | 5R |
| 8 | 5L | 2L | 5R | 2R |
| 9 | 5L | 2R | 2L | 5R |
| 10 | 5L | 2R | 5R | 2L |
| 11 | 5L | 5R | 2L | 2R |
| 12 | 5L | 5R | 2R | 2L |
| 13 | 2R | 2L | 5R | 5L |
| 14 | 2R | 2L | 5L | 5R |
| 15 | 2R | 5L | 2L | 5R |
| 16 | 2R | 5L | 5R | 2L |
| 17 | 2R | 5R | 2L | 5L |
| 18 | 2R | 5R | 5L | 2L |
| 19 | 2L | 2R | 5R | 5L |
| 20 | 2L | 2R | 5R | 5L |
| 21 | 2L | 5R | 2R | 5L |
| 22 | 2L | 5R | 5L | 2R |
| 23 | 2L | 5L | 2R | 5R |
| 24 | 2L | 5L | 5R | 2R |

Note: Rats received 4 trials / day. R = Clockwise, L = Counter-clockwise

**Table S5.** Balanced Latin Square Design Used for Testing Schedule

| Rat | Test 1 | Test 2 | Test 3 | Test 4 | Pattern |
| --- | --- | --- | --- | --- | --- |
| 1 | PS5 | PC5 | PS2 | PC2 | DCBA |
| 2 | PC5 | PS2 | PC2 | PS5 | CBAD |
| 3 | PS2 | PC2 | PS5 | PC5 | BADC |
| 4 | PC2 | PS5 | PC5 | PS2 | ADCB |
| 5 | PC2 | PS2 | PC5 | PS5 | ABCD |
| 6 | PS2 | PC5 | PS5 | PC2 | BCDA |
| 7 | PC5 | PS5 | PC2 | PS2 | CDAB |
| 8 | PS5 | PC2 | PS2 | PC5 | DABC |
| 9 | PS2 | PS5 | PC2 | PC5 | BDAC |
| 10 | PS5 | PC2 | PC5 | PS2 | DACB |
| 11 | PC2 | PC5 | PS2 | PS5 | ACDB |
| 12 | PC5 | PS2 | PS5 | PC2 | CBDA |
| 13 | PC5 | PC2 | PS5 | PS2 | CADB |
| 14 | PC2 | PS5 | PS2 | PC5 | ADBC |
| 15 | PS5 | PS2 | PC5 | PC2 | DBCA |
| 16 | PS2 | PC5 | PC2 | PS5 | BCAD |
| 17 | PC5 | PS5 | PS2 | PC2 | CDBA |
| 18 | PS5 | PS2 | PC2 | PC5 | DBAC |
| 19 | PS2 | PC2 | PC5 | PS5 | BACD |
| 20 | PC2 | PC5 | PS5 | PS2 | ACDB |

Note: PC2=A; PS2=B; PC5=C; PS5=D.
Infusions made “Pre-Choice” = PC, infusions made “Pre-Sample” = PS.
Animals were tested with an arm separation of 2 or 5.

**Table S6.** Experiment 1 (DNMP) design across the four test days

| Between-Subject | Within-Subject | Within-Subject | Within-Subject |
| --- | --- | --- | --- |
| Group | Infusion Time | Arm Separation | Trial |
| ISO | PS | S2 | HAB-BASE-TEST |
| ISO | PS | S5 | HAB-BASE-TEST |
| ISO | PC | S2 | HAB-BASE-TEST |
| ISO | PC | S5 | HAB-BASE-TEST |
| PRO | PS | S2 | HAB-BASE-TEST |
| PRO | PS | S5 | HAB-BASE-TEST |
| PRO | PC | S2 | HAB-BASE-TEST |
| PRO | PC | S5 | HAB-BASE-TEST |

Note: ISO = Isoproterenol; PRO = Propranolol; PS = infusions made Pre-Sample. PC = infusions made Pre-Choice. Animals were tested with an arm separation of 2 (S2) or 5 (S5). HAB = habituation trial; BASE = baseline trial; TEST = test trial.

**Table S7.** Cardinal direction rat was facing at the start of the trial.

| Direction | Number of Trials | Percentage of Trials (%) |
| --- | --- | --- |
| North | 236 | 27.76 |
| West | 227 | 26.71 |
| South | 191 | 22.47 |
| East | 196 | 23.06 |

Note: Total number of trials (not including habituation which was not videotaped) = 850.

## Notes

### Competing Interest Statement

The authors have declared no competing interest.

### Summary of Updates

Figures have been colorized, one figure has been added, one figure caption has been added, one figure has been revised, one figure has been removed, one table has been revised, three references have been added. The text has been amended to reflect these changes.

